# Quercetin treatments alleviate production, egg quality, blood metabolites, and anti-oxidant defence in laying hens : A meta-analysis study

**DOI:** 10.1101/2025.05.16.654414

**Authors:** Slamet Hartanto, Heru Ponco Wardono, Heri Kurnianto, Franciscus Rudi Prasetyo Hantoro, Amrih Prasetyo, Bambang Haryanto, Rini Nur Hayati, Dini Dwi Ludfiani, Rita Purwasih, Aan Andri Yano, Joko Sujiwo, Aera Jang, Sugiharto Sugiharto

**Affiliations:** Research Center for Sustainable Production System and Life Cycle Assessment, National Research and Innovation Agency, Serpong, Banten 15314, Indonesia; Research Center for Animal Husbandry, National Research and Innovation Agency, Bogor, West Java 16911, Indonesia; Research Center for Veterinary Science, National Research and Innovation Agency, Bogor, West Java 16915, Indonesia; Department of Agroindustry, Subang State Polytechnic, Subang, West Java 41285, Indonesia; Vocational School, Universitas Sebelas Maret, Surakarta, Central Java 57126, Indonesia; Department of Animal Production, Faculty of Animal Science, Universitas Gadjah Mada, Sleman, Special Region of Yogyakarta 55281, Indonesia; Department of Applied Animal Science, Kangwon National University, Chuncheon 24341, Korea; Faculty of Animal and Agricultural Sciences, Diponegoro University, Semarang, Central Java 50275, Indonesia

**Keywords:** Flavonoid, Egg, Layer, Supplementation, Systematic Statistical Analysis

## Abstract

**Objective:** The study aimed to evaluate impact of quercetin supplementation on performance, egg production, egg quality, blood metabolites, malondialdehyde (MDA) levels and anti-oxidant activity in laying hen

**Methods:** The meta-analysis study synthesized 27 included studies. We applied mean difference (MD) calculation using restricted maximum likelihood random-effects model to determine the effect size of quercetin treatments. A mixed-effects model was used in subgroup analysis to evaluate the influence of quercetin dose.

**Results:** Quercetin treatments increased (p<0.05) laying rate (LR) (MD = 2.819%), egg weight (EW) (MD = 1.209 g/unit), haugh unit (HU) (MD = 1.838%), shell thickness (ST) (MD = 0.014 mm), and yolk colour (YC) (MD = 0.526). Feed-to-egg ratio (FER) was reduced (MD = -0.146; p<0.05) in quercetin-treated laying hens. Quercetin decreased (p<0.05) serum glutamic pyruvic transaminase (SGPT) (MD = - 7.009 U/L), glucose (MD = -17.589 mg/dL), and total cholesterol (MD = -20.834) levels in laying hens. High-density lipoprotein (HDL) concentration was improved (MD = 32.590 mg/dL; p<0.05) by quercetin supplementation. Quercetin dietary reduced (MD = -7.373 nmol/mL; p<0.05) MDA level. Moreover, superoxide dismutase (SOD) concentration (MD = 8.114 U/mL; p<0.05) was enhanced in quercetin-administrated laying hens. Quercetin dose linearly affected (p<0.05) the SGPT, glucose, and total cholesterol levels. However, quercetin dose had quadratic effect (p<0.05) on LR, FER, and SOD content.

**Conclusion:** quercetin treatments effectively improved performance, egg production, egg quality, blood metabolites, and anti-oxidant defence in laying hens. The effective quercetin dose for optimizing LER, FER, and SOD content was 400 to 600 mg/kg.

## INTRODUCTION

Poultry contribute significantly to food security and nutrition by efficiently converting agri-food by-products into nutrient-rich meat and eggs through rapid production cycles [1]. However, they are susceptible to diseases and environmental stressors that lead to declines in both their quality and productivity [2]. The decrease of productivity is a vital threat to the sustainability of the poultry industry. Nutrition is a key strategy for enhancing productivity and reducing disease in poultry industry [3]. Natural bioactive compounds have crucial role in nutritional intervention to improve the productivity and health of laying hens [4].

Quercetin is a flavonoid renowned for its strong antioxidant, anti-inflammatory, and metabolic regulatory effects [5]. Quercetin dietary has been shown to reduce malondialdehyde (MDA) levels and enhance antioxidant defences in various animal studies [6]. In laying hens, the inclusion of quercetin has been studied as a method to enhance performance and egg quality. Quercetin supplementation modulates key performance indices in laying hens, such as feed intake (FI) [7–9], laying rate (LR) [10,11,12–15], and feed-to-egg ratio (FER) [16–18]. Similar findings are observed on haugh unit (HU) [16,17,19], shell thickness (ST) [16,20,21], egg weight (EW) [15,17,21], and yolk colour (YC) [17,18,20]. Furthermore, quercetin administrations lower MDA levels [21–23] and ameliorate antioxidant defences, as indicated by catalase (CAT) [12,14,21] and superoxide dismutase (SOD) [17,21,22] concentrations in laying hens. The significant impacts are found on serum glutamate pyruvate transaminase (SGPT) [10], glucose [10,12], total cholesterol [12,23,24], high-density lipoprotein (HDL) [12,24], and low-density lipoprotein (LDL) [12,24] in quercetin-treated hens. Additionally, quercetin treatments improve albumen quality [25], Ca deposition in reproductive organs [26], and hepatoprotective activities [27] in laying hens.

However, contradictory findings are demonstrated by certain studies. Quercetin has no effect on FI [28–30], LR [21,31,32], and FER [23,28,31] in laying hens. Similarly, no influences are detected in EW [14,29,32], HU [28,31,33], ST [28,31,33], and YC [24,28,33] of quercetin-administrated hens. Quercetin supplementations do not affect blood metabolites including glucose [18,19], total cholesterol [8,18,22], HDL [18,19], and LDL [18]. Moreover, SOD [18] and CAT [17,22] levels in hens remain unaffected by quercetin treatments. These inconsistencies results are attributed to differences in quercetin dose, duration of treatment, and age of hens. A systematic statistical study is essential to conclude the precise dose, duration, and age of hens for quercetin treatment in laying hens.

Meta-analysis is a method to systematically analyze data from previous contradictory findings to obtain robust conclusions [34,35]. The study aimed to compute multiple published data on performance traits, egg quality, blood metabolites, MDA levels, and antioxidant defence status in quercetin-supplemented laying hens using meta-analysis approach.

## MATERIALS AND METHODS

### Protocol for Searching and Selecting Included Studies

The guidelines of the Preferred Reporting Items for Systematic Reviews and Meta-Analyses (PRISMA) were employed as the protocol for our study (Fig 1). Initially, we performed a rigorous literature search through Scopus and Web of Science, using a combination of keywords as shown in Table 1. The titles and abstracts were reviewed to screen the eligible study by 5 reviewers (SH, HK, HPW, FRPH, AP, and BH). The duplicated article were merged in this step. Moreover, two reviewers (SH and HK) independently assessed the full-text articles for the included studies. A third-party reviewer (HPW) was involved to resolve any disagreements. The established criteria for included studies as follows: (1) Original research studies; (2) Articles published in English; (3) The full text were available; (4) Studies on quercetin supplementation in laying hens; (5) Studies on effect of quercetin on production, egg quality, antioxidant status, and blood metabolite parameters; (6) a control group in the studies was included; and (7) measures of variability (confidence intervals [CI], standard errors [SE], or standard deviations [SD]) was reported. Additionally, a reviewer (SH) scrutinized the references of the eligible studies to identify any missed article. All included studies was analysed using the Systematic Review Centre for Laboratory Animal Experimentation’s (SYRCLE’s) tool to determine the risk of bias.

**Figure 1.**
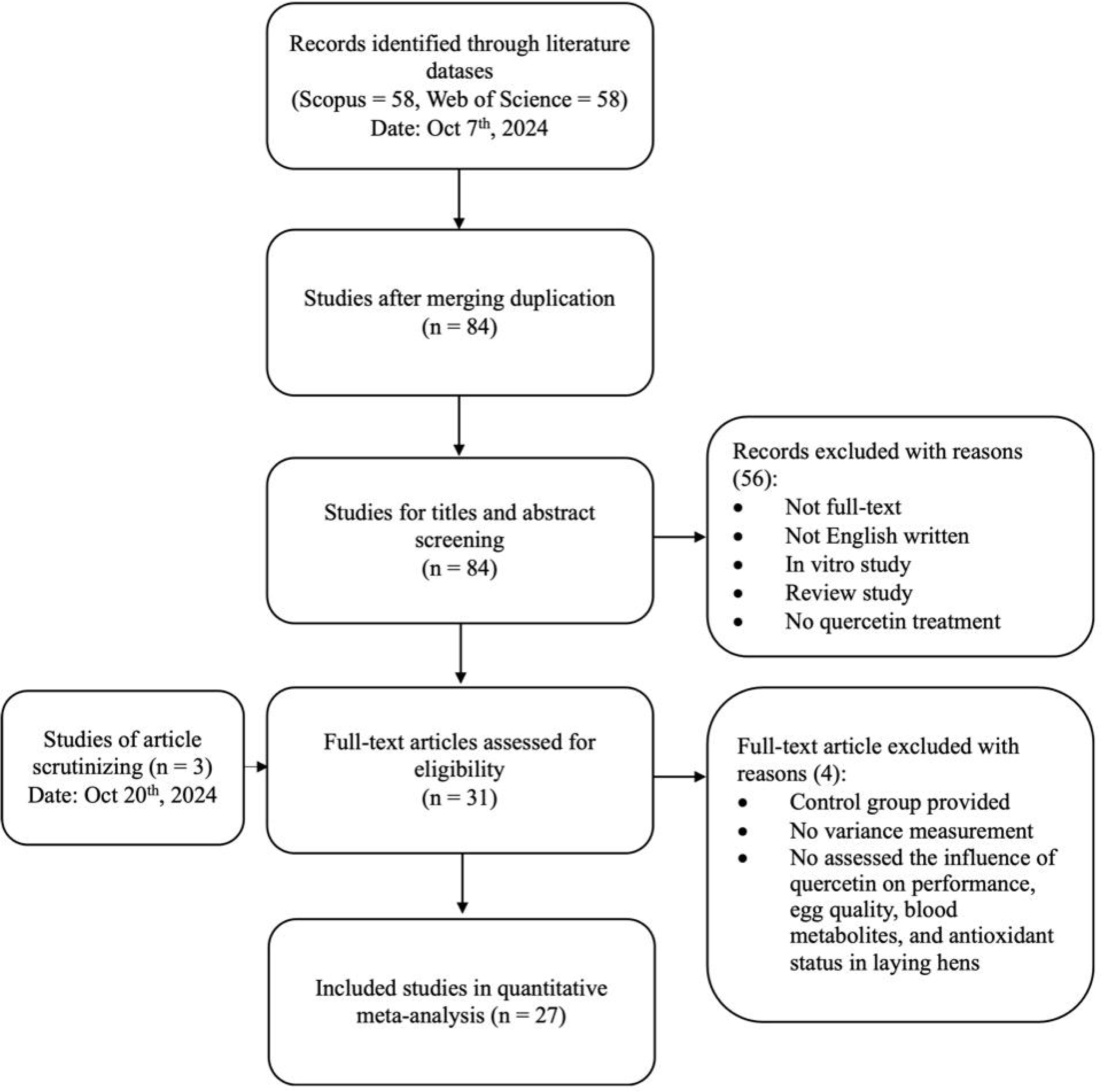
PRISMA workflow for literature strategy

**Figure 2.**
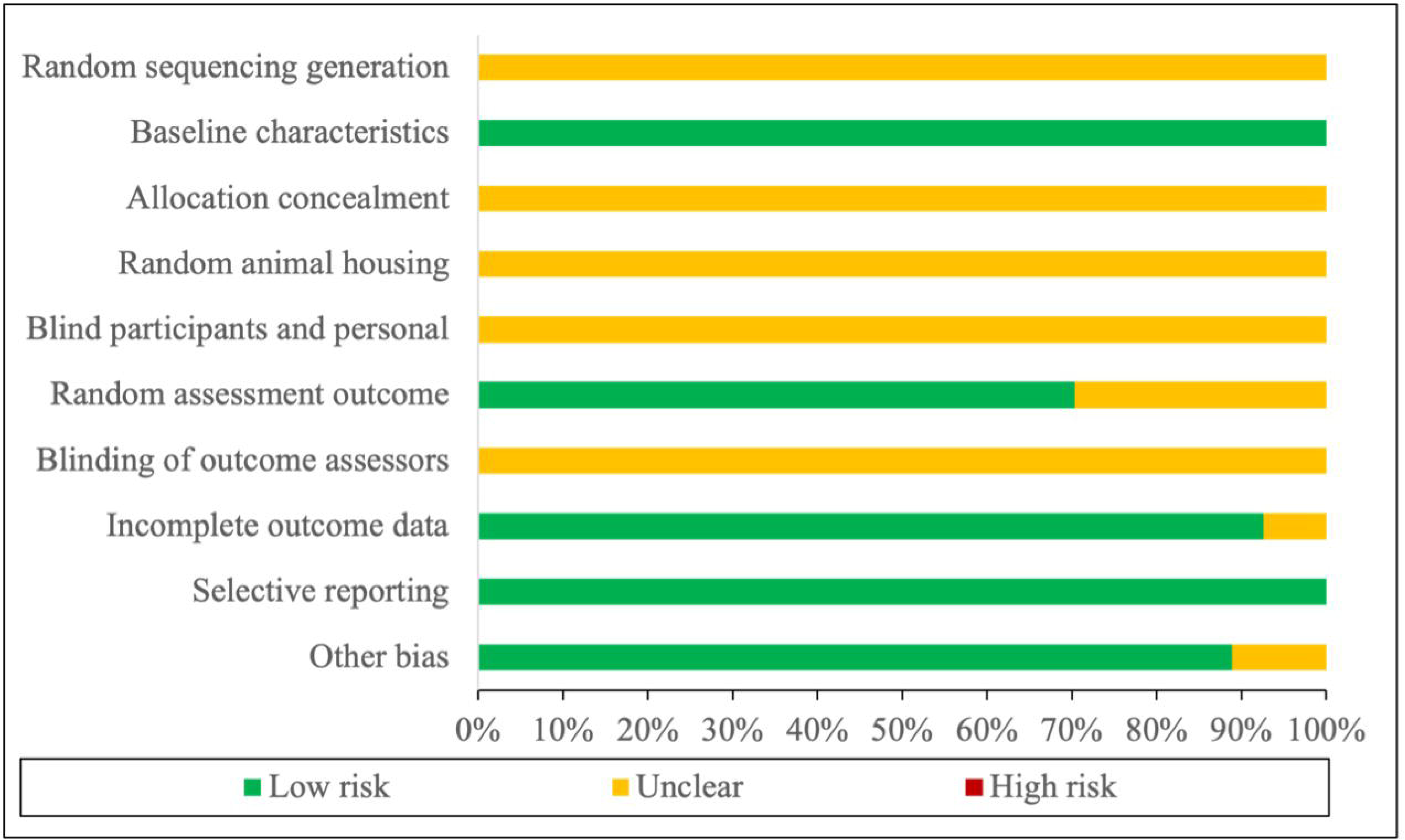
SYRCLE risk of bias

**Table 1.** Keywords combinations for literature search.

| Databases | Keywords | Studies |
| --- | --- | --- |
| Scopus | ( TITLE-ABS-KEY ( hen* ) AND TITLE-ABS-KEY ( quercetin ) AND<br>TITLE-ABS-KEY ( egg* ) ) | 58 |
| Web of<br>Science | #1 TS=(hen*)<br>#2 TS=(quercetin)<br>#3 TS=(egg*)<br>#1 AND #2 AND #3 | 58 |

### Data Acquisition Process

The characteristics of included study are detailed in Table 2. The data extracted in the study were key performance indicators (FI and FER), egg production (LR and EW), egg quality parameters (ST, YC, EW, and HU), blood metabolites characteristics (SGPT, glucose, cholesterol, HDL, and LDL contents), oxidative stress indicators (MDA level), and antioxidant enzyme activities (CAT and SOD levels).

**Table 2.**
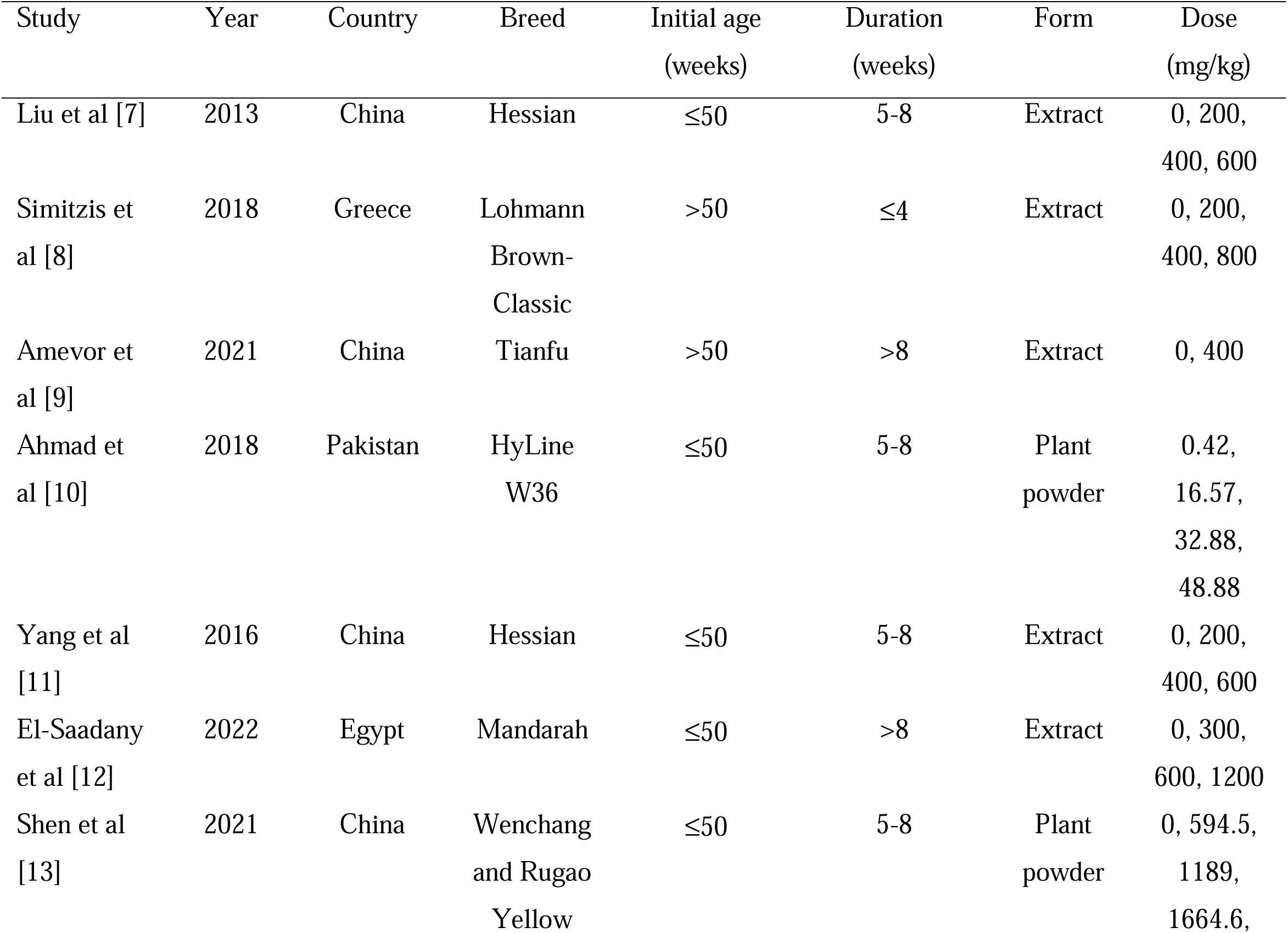
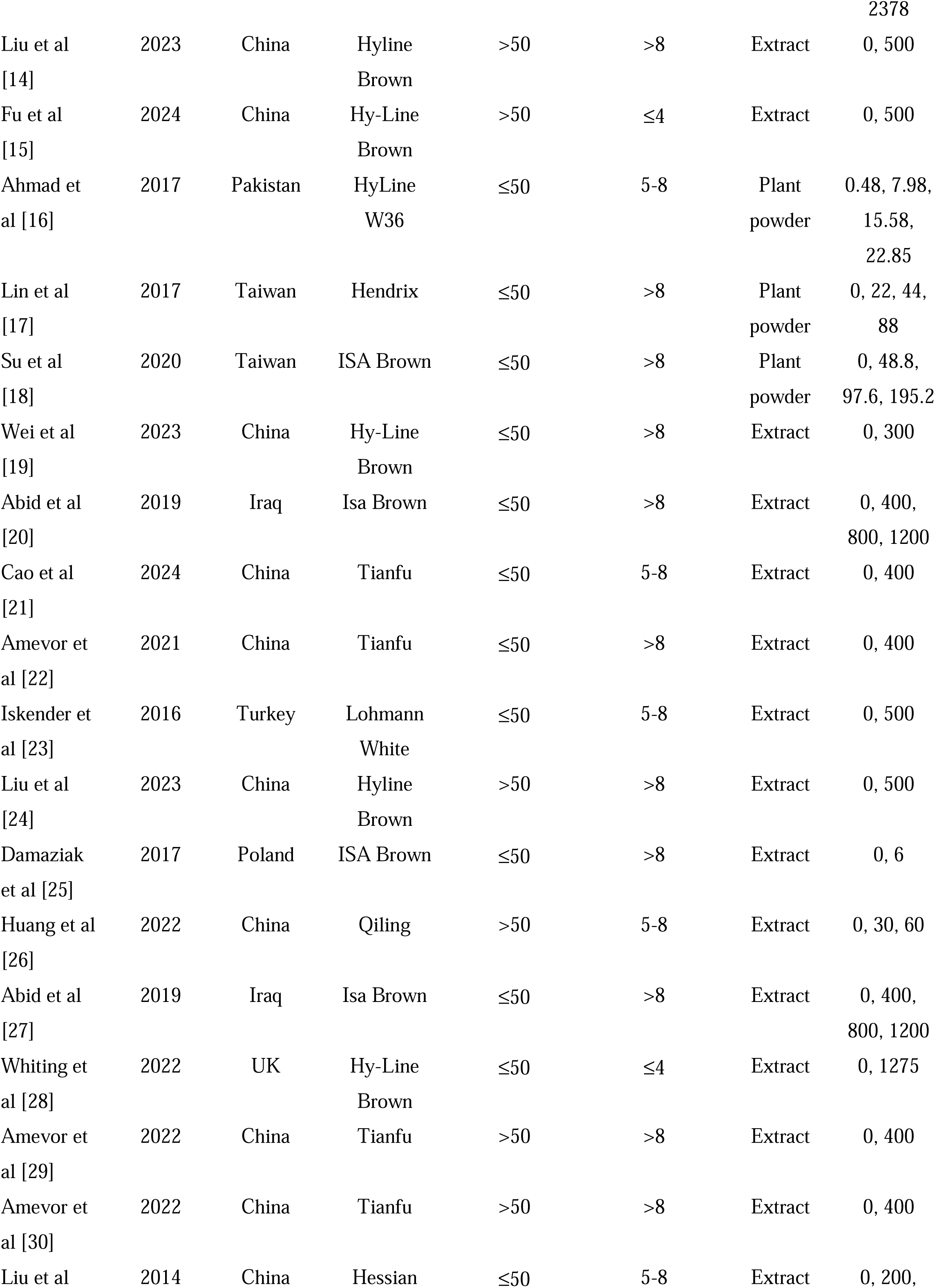
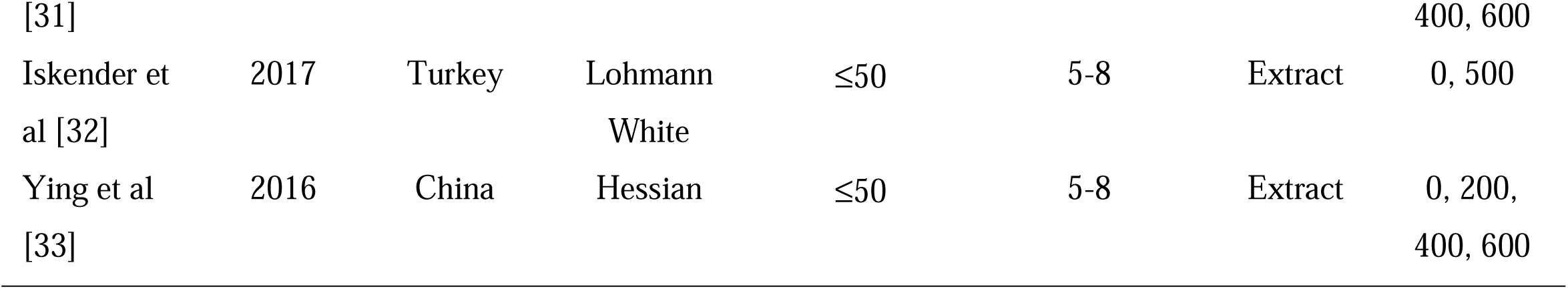
Included studies attributions.

Mean and SD were extracted from included studies to perform the meta-analysis. The SD was computed using two methods: (1) SD = SE√N, where N denotes the number of replicates; and (2) SD = √N × (upper CI – lower CI)/3.92, where 3.92 corresponds to the standard error for a 95% confidence interval. For small sample sizes (<60), the constant 3.92 was replaced with the appropriate t-distribution value based on the degrees of freedom [36]. Graphical data were digitized using WebPlotDigitizer software [37].

### Statistical analysis

The study calculated mean difference (MD) using restricted maximum likelihood (REML) random-effects model was applied to evaluate effect sizes in their original units [38]. The *I²* statistic was applied to quantify the degree of heterogeneity, with values >50%, 25-50%, and <25% indicating high, moderate, and low heterogeneity, respectively [39]. To assess publication bias, a funnel plot was constructed, and Egger’s regression test was employed to detect asymmetry [40]. The *p* value of egger’s test less than 0.05 implied that the study has a high risk publication bias.

Subgroup analyses were conducted when the following criteria were fulfilled: (1) the significancy of effect size was found (p<0.05), (2) significant heterogeneity was observed (I² > 50%, p<0.001), and (3) number of studies was sufficient (≥ 10 comparisons) [41]. Quercetin dose was classified as a continuous covariate. Meanwhile, the initial age, treatment duration, and quercetin form were categorical covariates. To assess the impact of quercetin dose, we applied a mixed-effects model for subgroup analysis. The equation was presented as follows:

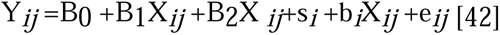

In this equation, Yij represents the dependent variable (where *i* indicates the study from 1 to *m*, and *j* the observation from 1 to *ni*). *B*_0_ denotes the overall intercept across all studies. *B*_1_ and *B*_2_ are the linear and quadratic coefficients of the explanatory variables, respectively. The predictor variable value (quercetin dose) is indicated by *Xij*. Moreover, *si* captures the random effect of the *i*-th study, *bi* is the random slope for the *i*-th study, and *eij* signifies the residual error not explained.

All analyses was conducted in R program [43] using using the “metafor” package [44]. Additionally, the results of subgroup analysis were visualized using the “ggplot2” package [45] and Microsoft excel [46].

## RESULTS

### Characteristics of included study

We included 27 eligible studies in the present study. The quercetin dose was 0 – 2,378 mg/kg. The initial age of less than 50 weeks and more than 50 weeks were 70.38% and 29.62%, respectively. Treatment duration consisted of 11.11% less than equal to 4 weeks, 40.74% 5 to 8 weeks, and 48.15% more than 8 weeks. The quercetin forms were 81.48% extract powder and 18.52% plant powder.

### Effect of quercetin dietary on production and egg quality

Administrations of quercetin affected (p<0.001) performance, production, and egg quality parameters in laying hens, including LR, FER, EW, HU, ST, and YC (Table 3). In contrast, quercetin supplementation had no effect (p> 0.05) on FI. Quercetin treatments increased LR (MD = 2.819%), egg weight (MD = 1.209 g/unit), HU (MD = 1.838%), ST (MD = 0.014 mm), and YC (MD = 0.526). Moreover, FER was reduced (MD = -0.146) in quercetin-treated laying hens.

**Table 3.** Effect of dietary quercetin on production and egg quality.

| Parameter | N | Control mean<br>(SD) | MD (95% CI) | p-value | Heterogeneity test |  |
| --- | --- | --- | --- | --- | --- | --- |
|  |  |  |  |  | p-value | I<sup>2</sup> (%) |
| Feed intake, g/d/hen | 42 | 99.488(4.856) | 0.902(-0.068; 1.872) | 0.068 | < .001 | 99.72 |
| Laying rate, % | 68 | 74.312(5.393) | 2.819(1.574; 4.064) | <.001 | < .001 | 99.46 |
| Feed-egg-ratio | 60 | 2.167(0.124) | -0.146(-0.198; -0.094) | <.001 | < .001 | 99.87 |
| Egg weight, g/unit | 70 | 60.214(3.141) | 1.209(0.702; 1.716) | <.001 | < .001 | 97.75 |
| Haugh unit, % | 65 | 82.298(7.580) | 1.838(0.901; 2.776) | <.001 | < .001 | 93.55 |
| Shell thickness, mm | 71 | 0.376(0.075) | 0.014(0.007; 0.021) | <.001 | < .001 | 89.33 |
| Yolk colour | 52 | 7.964(2.222) | 0.526(0.334; 0.717) | <.001 | < .001 | 92.95 |

N, number of comparisons; SD, standard deviation; MD, mean difference; I<sup>2</sup>, Inconsistency index.

### Effect of quercetin dietary on blood metabolites

Table 4 shows effect of quercetin dietary on blood metabolites. Glucose, SGPT, HDL, and total cholesterol levels were influenced (p<0.001) by quercetin dietary. However, LDL concentration in control and treated laying hens was not different (p<0.05). Quercetin effectively decreased SGPT (MD = -7.009 U/L), glucose (MD = -17.589 mg/dL), and total cholesterol (MD = -20.834) levels in laying hens. HDL concentration was improved (MD = 32.590 mg/dL) by quercetin supplementation.

**Table 4.** Blood metabolite parameters in control and quercetin treated laying hens.

| Parameter | N | Control mean (SD) | MD (95% CI) | p-value | Heterogeneity test |  |
| --- | --- | --- | --- | --- | --- | --- |
|  |  |  |  |  | p-value | I<sup>2</sup> (%) |
| SGPT, U/L | 15 | 20.318(1.210) | -7.009(-9.196; -4.822) | <.001 | < .001 | 98.95 |
| Glucose, mg/dL | 16 | 248.060(8.281) | -17.589(-23.627; -11.550) | <.001 | < .001 | 96.55 |
| Total cholesterol, mg/dL | 30 | 150.287(12.529) | -20.834(-38.721; -2.948) | 0.022 | < .001 | 99.81 |
| HDL, mg/dL | 14 | 52.068(8.270) | 32.590(2.257; 62.924) | 0.035 | < .001 | 99.73 |
| LDL, mg/dL | 14 | 59.593(9.478) | -8.552(-17.146; 0.042) | 0.051 | < .001 | 95.31 |

N, number of comparisons; SD, standard deviation; MD, mean difference; I<sup>2</sup>, inconsistency index; SGPT, serum glutamate pyruvate transaminase; HDL, high-density lipoprotein; LDL, low-density lipoprotein.

### Impact of quercetin supplementation on MDA level and antioxidant activity

Quercetin treatment influenced (p<0.001) MDA and SOD levels in laying hens (Table 5). On the contrary, significant effect was not observed (p> 0.05) on CAT concentration. Quercetin dietary reduced (MD = - 7.373 nmol/mL) MDA level. Contrary, SOD concentration (MD = 8.114 U/mL) was enhanced in quercetin-administrated laying hens.

**Table 5.** Comparison of MDA level and antioxidant status between control and quercetin treatment in laying hens.

| Parameter | N | Control mean<br>(SD) | MD (95% CI) | p-value | Heterogeneity test |  |
| --- | --- | --- | --- | --- | --- | --- |
|  |  |  |  |  | p-value | I<sup>2</sup> (%) |
| MDA, | 16 | 19.436(2.586) | -7.373(-12.304; -2.443) | 0.003 | < .001 | 99.97 |
| nmol/mL |  |  |  |  |  |  |
| SOD, U/mL | 19 | 39.722(3.018) | 8.114(0.233; 15.995) | 0.044 | < .001 | 100 |
| CAT, U/mL | 15 | 23.363(7.976) | 3.790(-0.521; 8.102) | 0.085 | < .001 | 97.6 |

N, number of comparisons; SD, standard deviation; MD, mean difference; $I^2$ , inconsistency index; MDA, malondialdehyde; SOD, superoxide dismutase; CAT, catalase.

### Heterogeneity and subgroup analyses

Heterogeneity of all studies was significant (p<0.001) and high (*I_2_* > 50%) (Table 3 – 5). Furthermore, subgroup analysis found that quercetin dose had effects (p<0.05) on LR (R^2^ = 11.44%), FER (R^2^ = 9.62%), SGPT (R^2^ = 33.07%), glucose (R^2^ = 29.42%), cholesterol (R^2^ = 28.01%), and SOD (R^2^ = 30.71%) (Table 6). Treatment duration influenced (p<0.05) LR (R^2^ = 29.42%), YC (R^2^ = 29.42%), SGPT (R^2^ = 29.42%), glucose (R^2^ = 29.42%), and cholesterol (R^2^ = 29.42%). Moreover, initial age of hens impacted (p<0.05) LR (R^2^ = 5.18%), ST (R^2^ = 24.58%), and SOD (R^2^ = 24.89%). Also, quercetin forms affected (p<0.05) LR (R^2^ = 4.99%), FER (R^2^ = 9.75%), YC (R^2^ = 16.98%), cholesterol (R^2^ = 26.17%), and SOD (R^2^ = 22.11%).

**Table 6.** Moderator test of meta-regression of quercetin treatment in laying hens.

| Parameter | Covariates | QM | df | *p*-value | R<sup>2</sup> (%) |
| --- | --- | --- | --- | --- | --- |
| Laying rate | Dose | 9.602 | 2 | 0.008 | 11.44 |
|  | Duration | 1.252 | 2 | 0.535 | 0 |
|  | Initial age | 3.905 | 1 | 0.048 | 5.18 |
|  | Form | 4.138 | 1 | 0.042 | 4.99 |
| Feed-to-egg ratio | Dose | 7.399 | 2 | 0.025 | 9.62 |
|  | Duration | 2.269 | 2 | 0.322 | 0 |
|  | Initial age | 0.3 | 1 | 0.584 | 0 |
|  | Form | 5.9 | 1 | 0.015 | 9.75 |
| Egg weight | Dose | 2.293 | 1 | 0.130 | 2.72 |
|  | Duration | 1.575 | 2 | 0.455 | 0.29 |
|  | Initial age | 2.162 | 1 | 0.141 | 4.74 |
|  | Form | 0.481 | 1 | 0.488 | 0 |
| Haugh unit | Dose | 2.848 | 1 | 0.091 | 2.48 |
|  | Duration | 3.454 | 2 | 0.178 | 0 |
|  | Initial age | 0.067 | 1 | 0.795 | 0 |
|  | Form | 0.012 | 1 | 0.914 | 0 |
| Shell thickness | Dose | 0.078 | 1 | 0.780 | 0 |
|  | Duration | 2.316 | 2 | 0.314 | 0.29 |
|  | Initial age | 18.014 | 1 | < .001 | 24.58 |
|  | Form | 1.691 | 1 | 0.193 | 1.78 |
| Yolk colour | Dose | 1.565 | 1 | 0.211 | 2.79 |
|  | Duration | 6.595 | 2 | 0.037 | 9.33 |
|  | Initial age | 0.075 | 1 | 0.784 | 0 |
|  | Form | 6.936 | 1 | 0.008 | 16.98 |
| SGPT | Dose | 7.741 | 1 | 0.005 | 33.07 |
|  | Duration | 42.522 | 2 | < .001 | 75.29 |
|  | Initial age | NA | NA | NA | NA |
|  | Form | NA | NA | NA | NA |
| Glucose | Dose | 4.995 | 1 | 0.025 | 29.42 |
|  | Duration | 20.545 | 2 | < .001 | 60.63 |
|  | Initial age | NA | NA | NA | NA |
|  | Form | 0 | 1 | 0.994 | 0 |
| Total cholesterol | Dose | 11.635 | 1 | < .001 | 28.01 |
|  | Duration | 22.999 | 2 | < .001 | 44.79 |
|  | Initial age | 0.038 | 1 | 0.846 | 0 |
|  | Form | 10.426 | 1 | 0.001 | 26.17 |
| HDL | Dose | 0.632 | 1 | 0.426 | 0 |
|  | Duration | 0.003 | 1 | 0.959 | 0 |
|  | Initial age | 0.003 | 1 | 0.959 | 0 |
|  | Form | 1.117 | 1 | 0.291 | 0.87 |
| MDA | Dose | 0.034 | 1 | 0.853 | 0 |
|  | Duration | 1.295 | 2 | 0.523 | 0 |
|  | Initial age | 0.449 | 1 | 0.503 | 0 |
|  | Form | 0.621 | 1 | 0.431 | 0 |
| SOD | Dose | 8.706 | 2 | 0.013 | 30.71 |
|  | Duration | 0.605 | 2 | 0.739 | 0 |
|  | Initial age | 8.989 | 1 | 0.003 | 24.89 |
|  | Form | 5.328 | 1 | 0.021 | 22.11 |

QM, Coefficient of moderators; df, degree of freedom; $R^2$ , heterogeneity accounted for by covariate; NA, non- available; SGPT, serum glutamate pyruvate transaminase; HDL, high-density lipoprotein; MDA, malondialdehyde; SOD, superoxide dismutase.

Quercetin dose linearly influenced the SGPT, glucose, and total cholesterol levels (Table 7; Figures 3A-C). However, LR, FER, and SOD content were influenced quadratically by quercetin dose. The highest LR was found in the quercetin dose of 600 mg/kg (Fig 3D). The optimum FER was observed in the 400 to 600 mg/kg quercetin dose (Figure 3E). The treatment of 500 mg/kg modulated the highest SOD concentration (Figure 3F).

**Figure 3.**
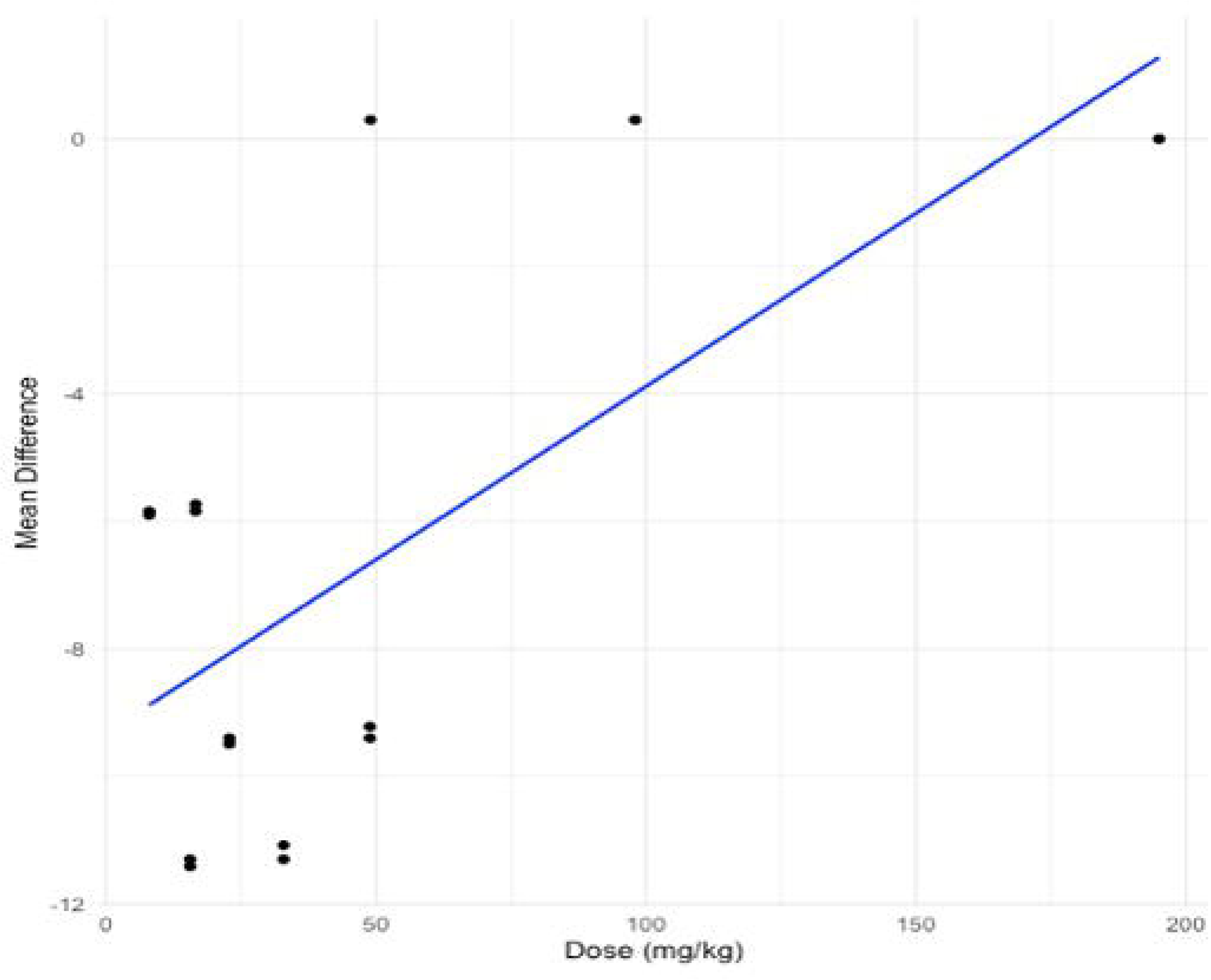
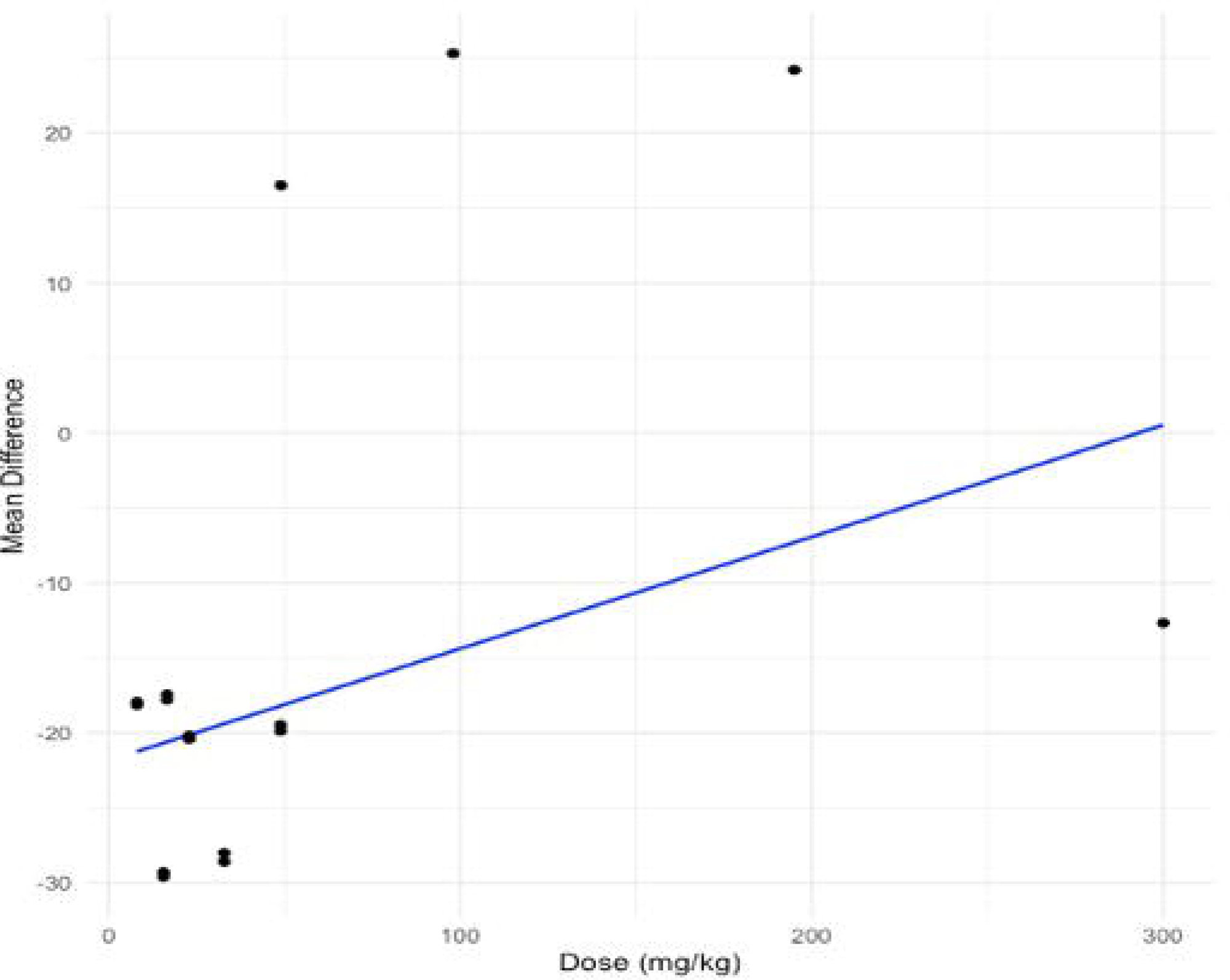
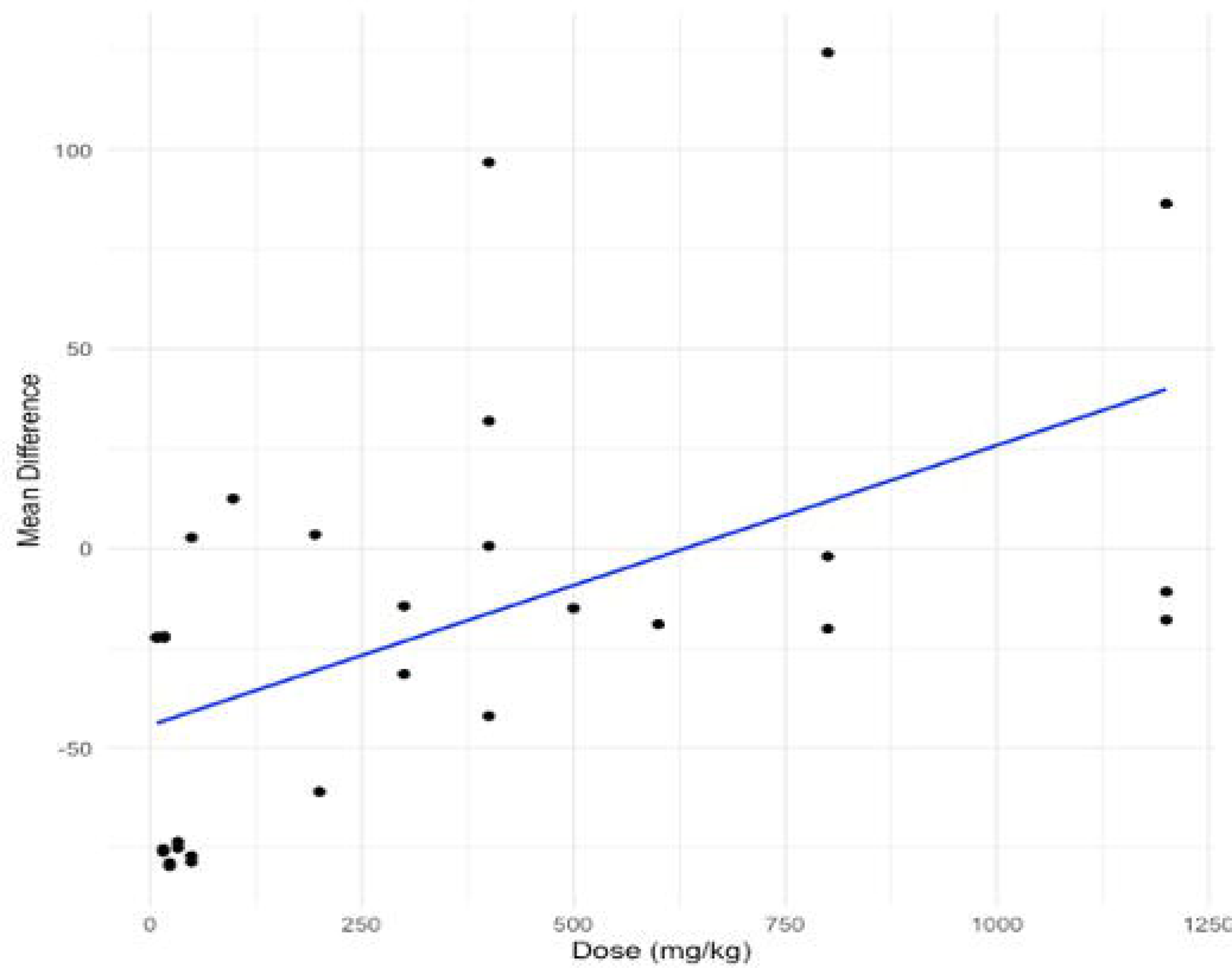
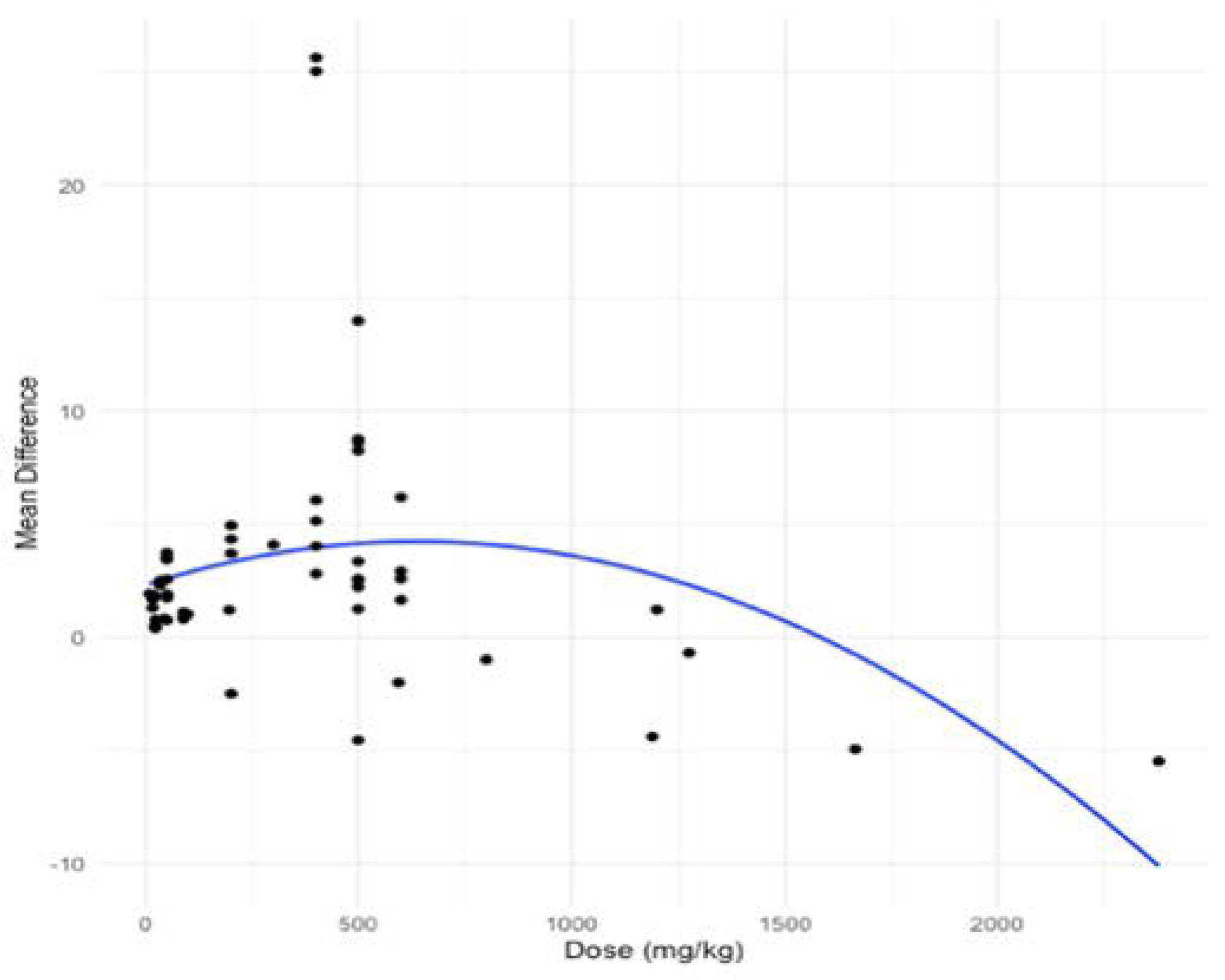
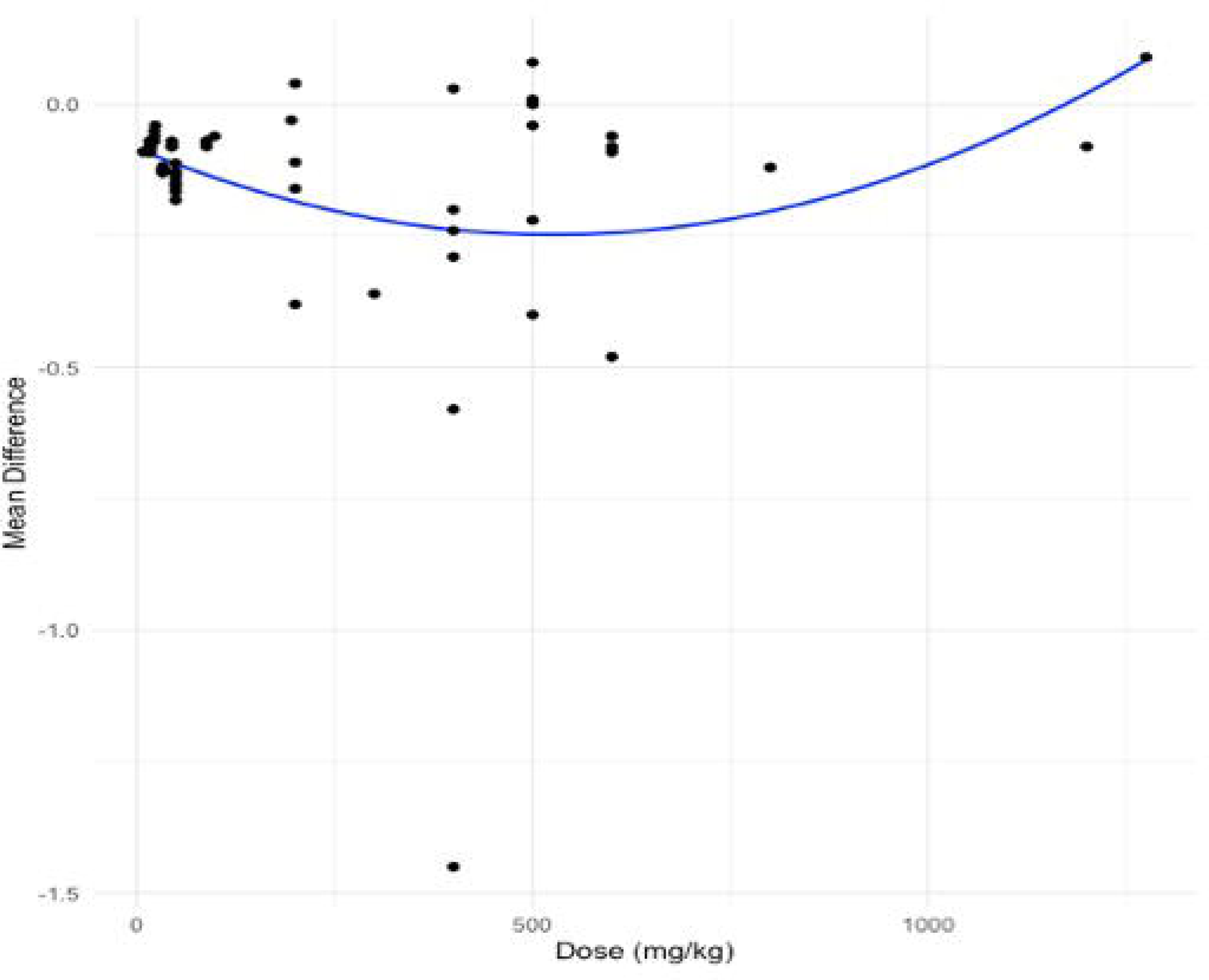
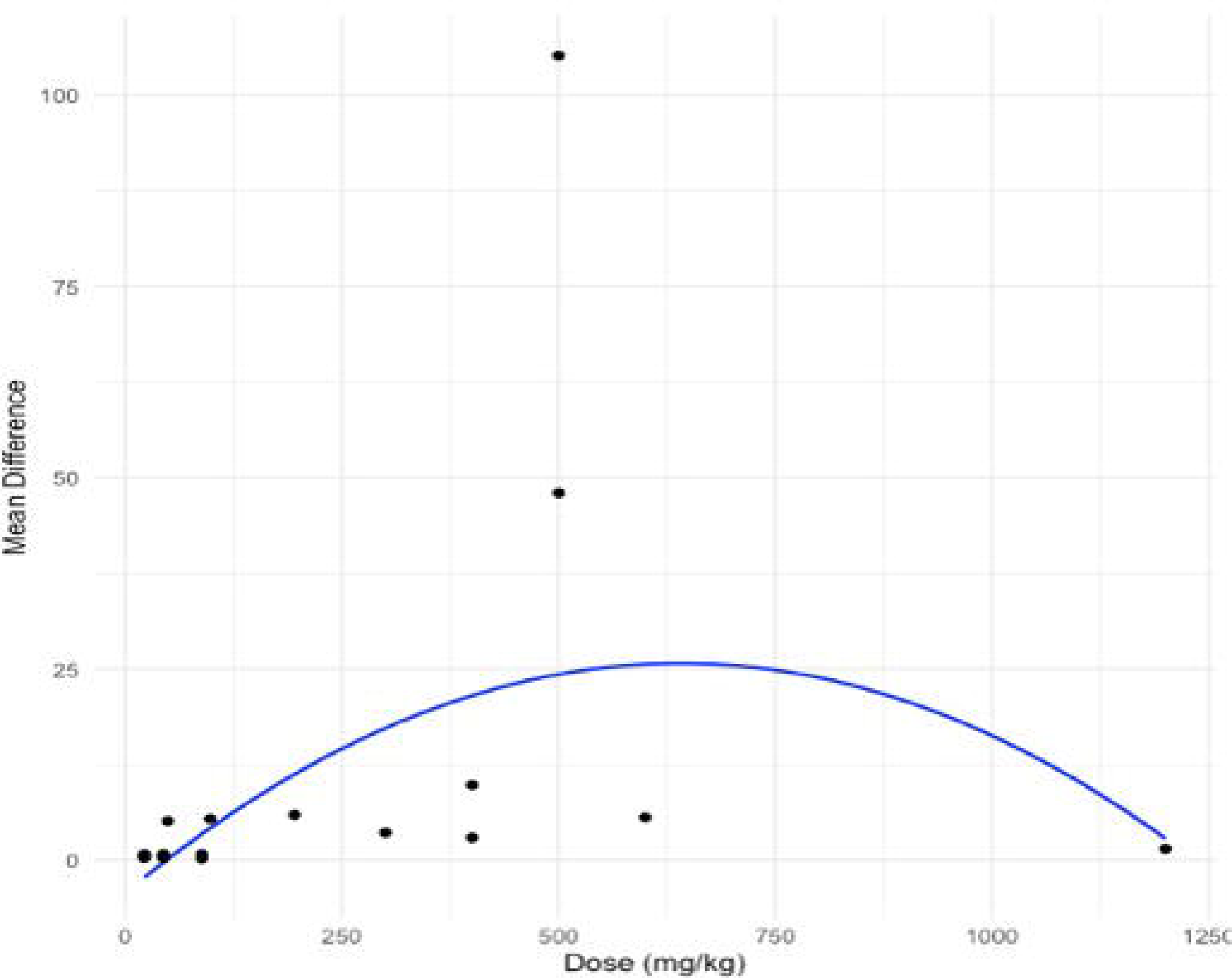
Subgroup analysis for effect of quercetin dose on, (a) SGPT (U/L), (b) glucose (mg/ dL), (c) cholesterol (mg/ dL), and (d) laying rate (%), (e) feed-to-egg ratio, (f) SOD (U/mL) in laying hens

**Table 7.**
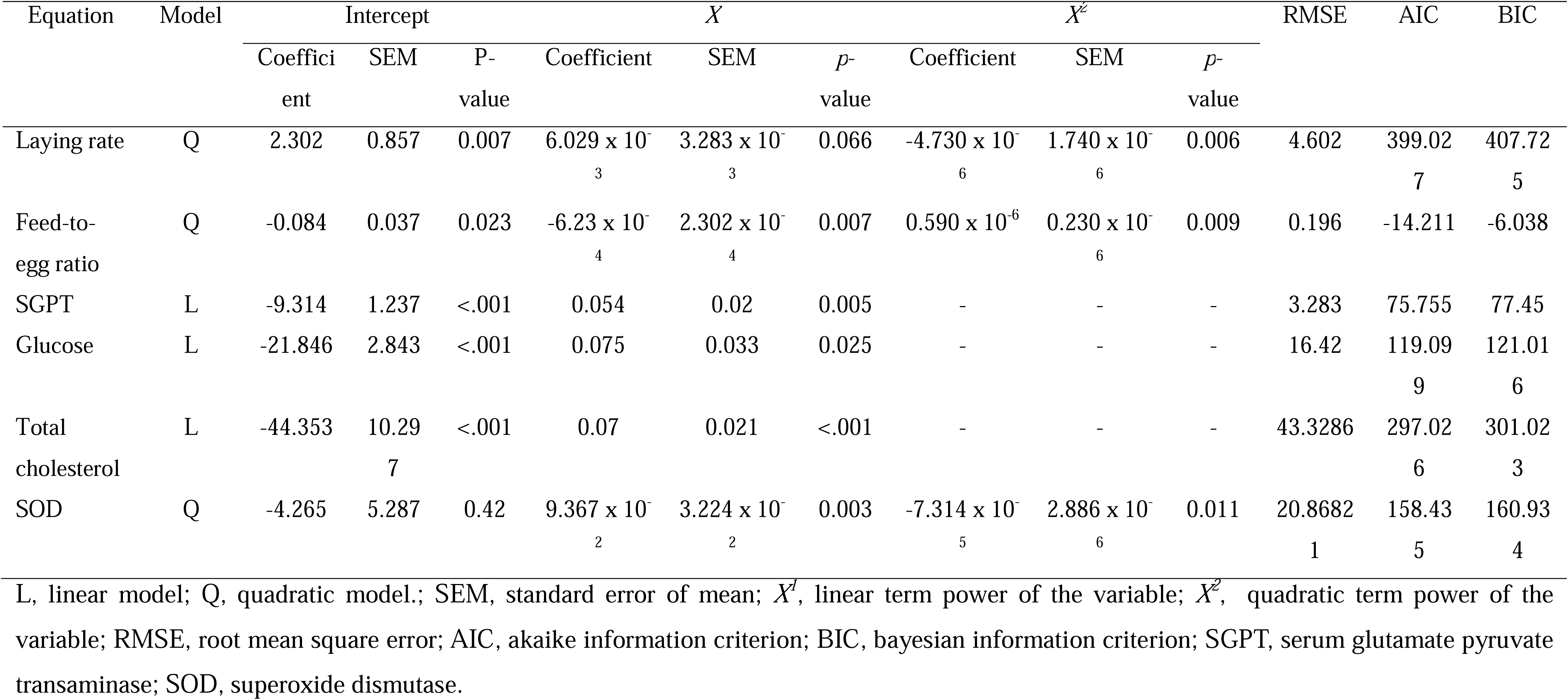
Subgroup analysis: regression model for relationship of quercetin dose on parameters in laying hens.

Treatment duration for 5 to 8 and more than 8 weeks enhanced (p<0.05) the YC (Figure 4A). In contrast, treatment for less than 4 weeks had no (p> 0.05) effect on YC. Figures 4B to 4D show that less than 4 and 5 to 8 weeks supplementations decreased (p<0.001) SGPT, glucose, and total cholesterol levels. However, the effect on SGPT, glucose, and cholesterol concentrations was not found in more than 8 weeks treatment.

**Figure 4.**
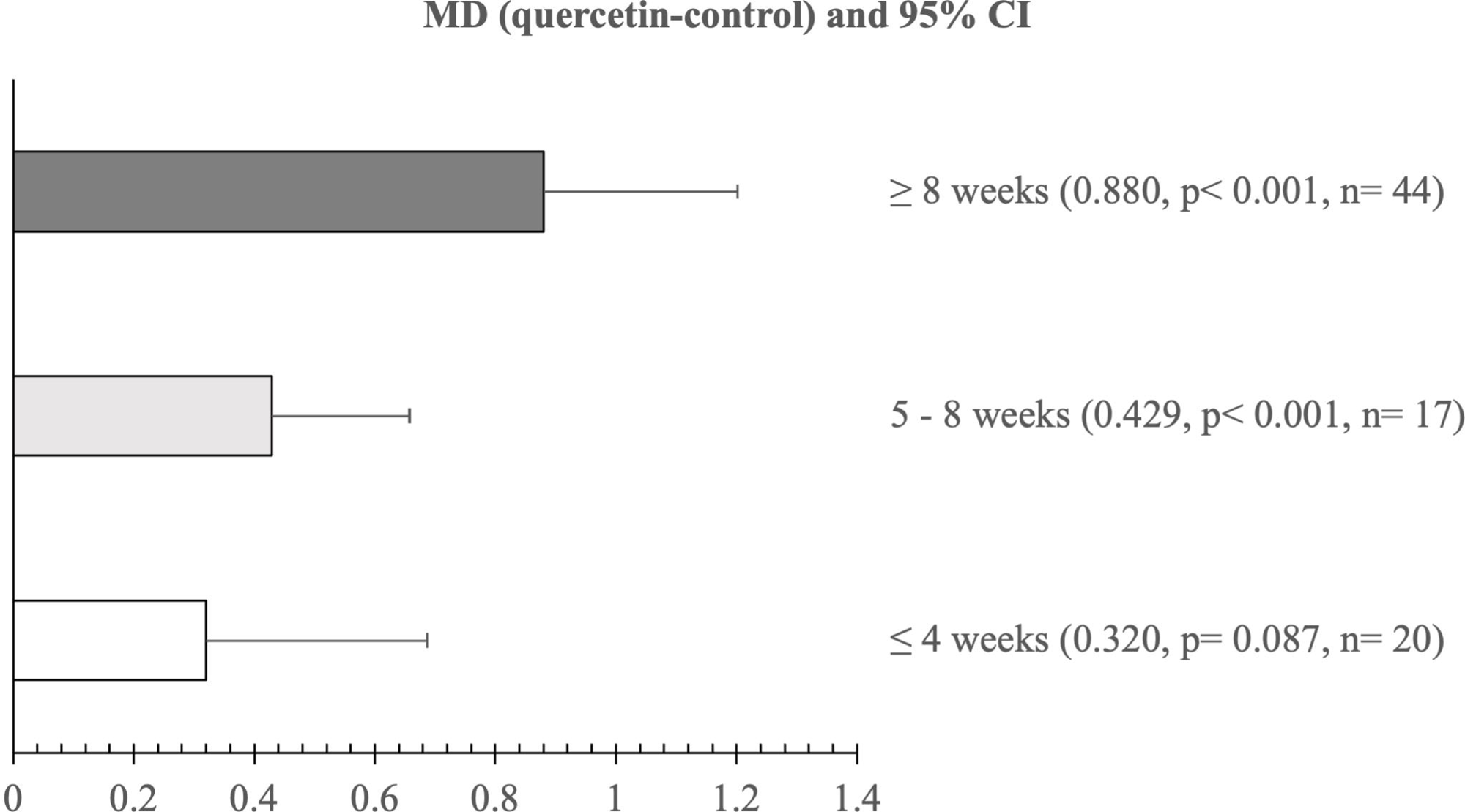
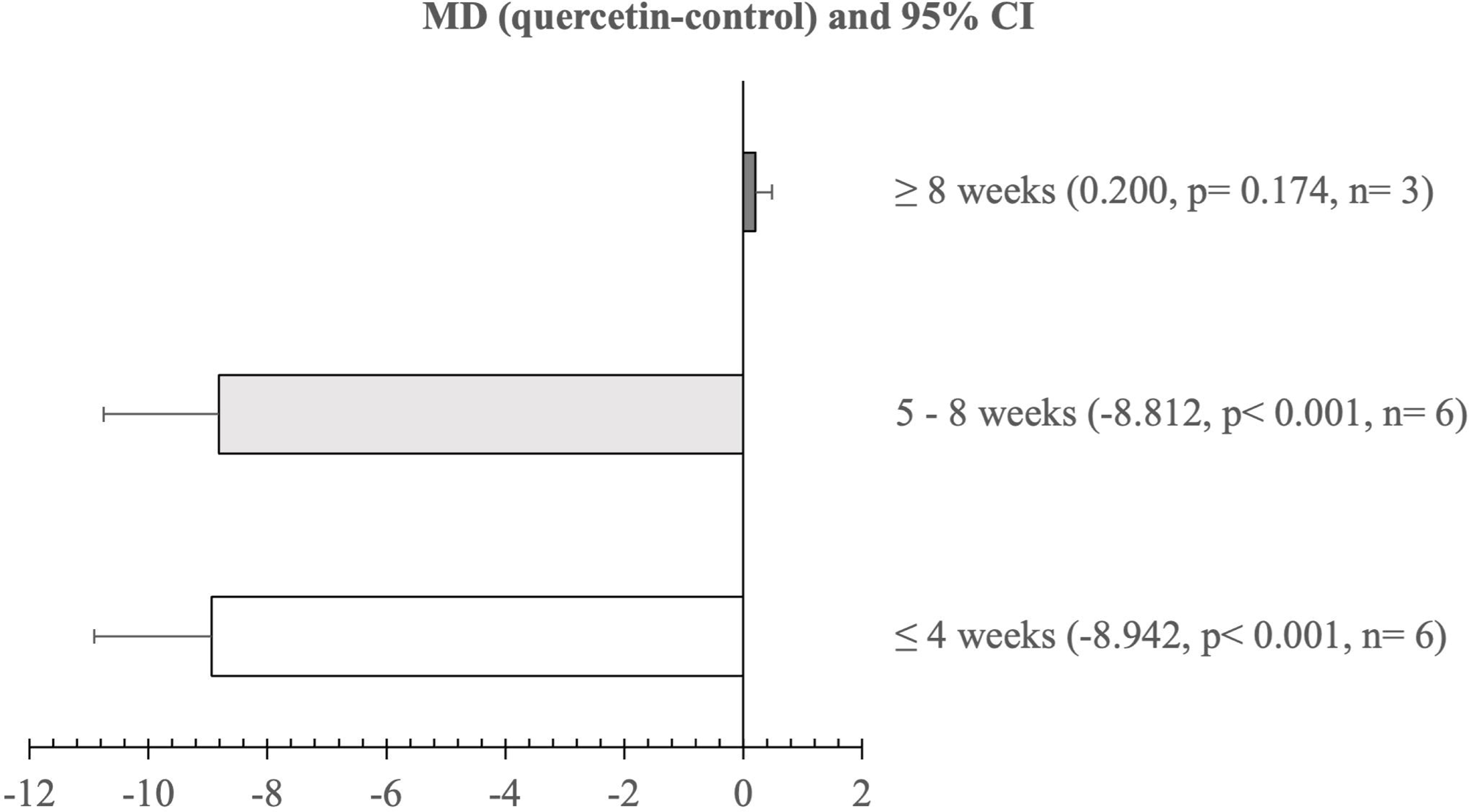
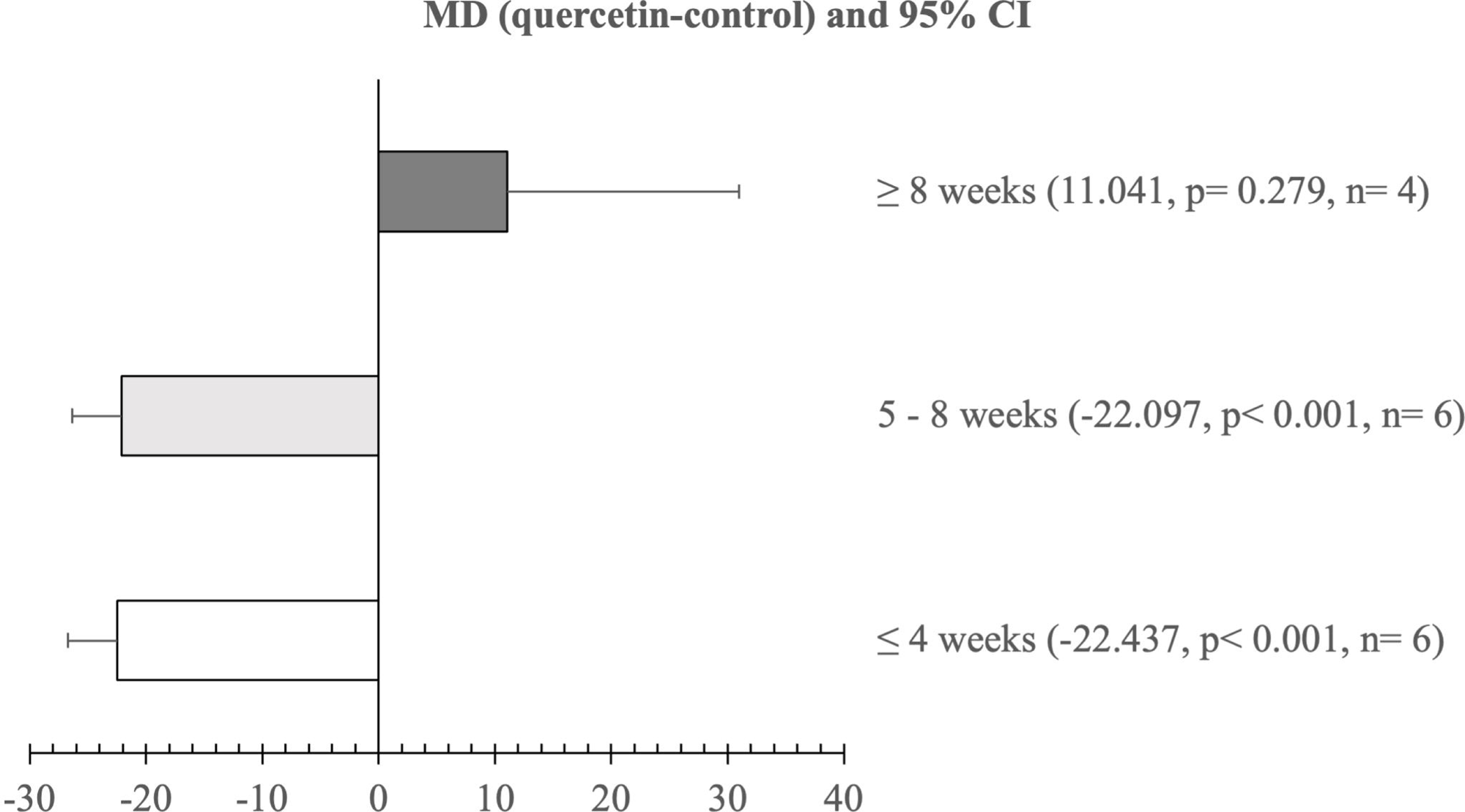
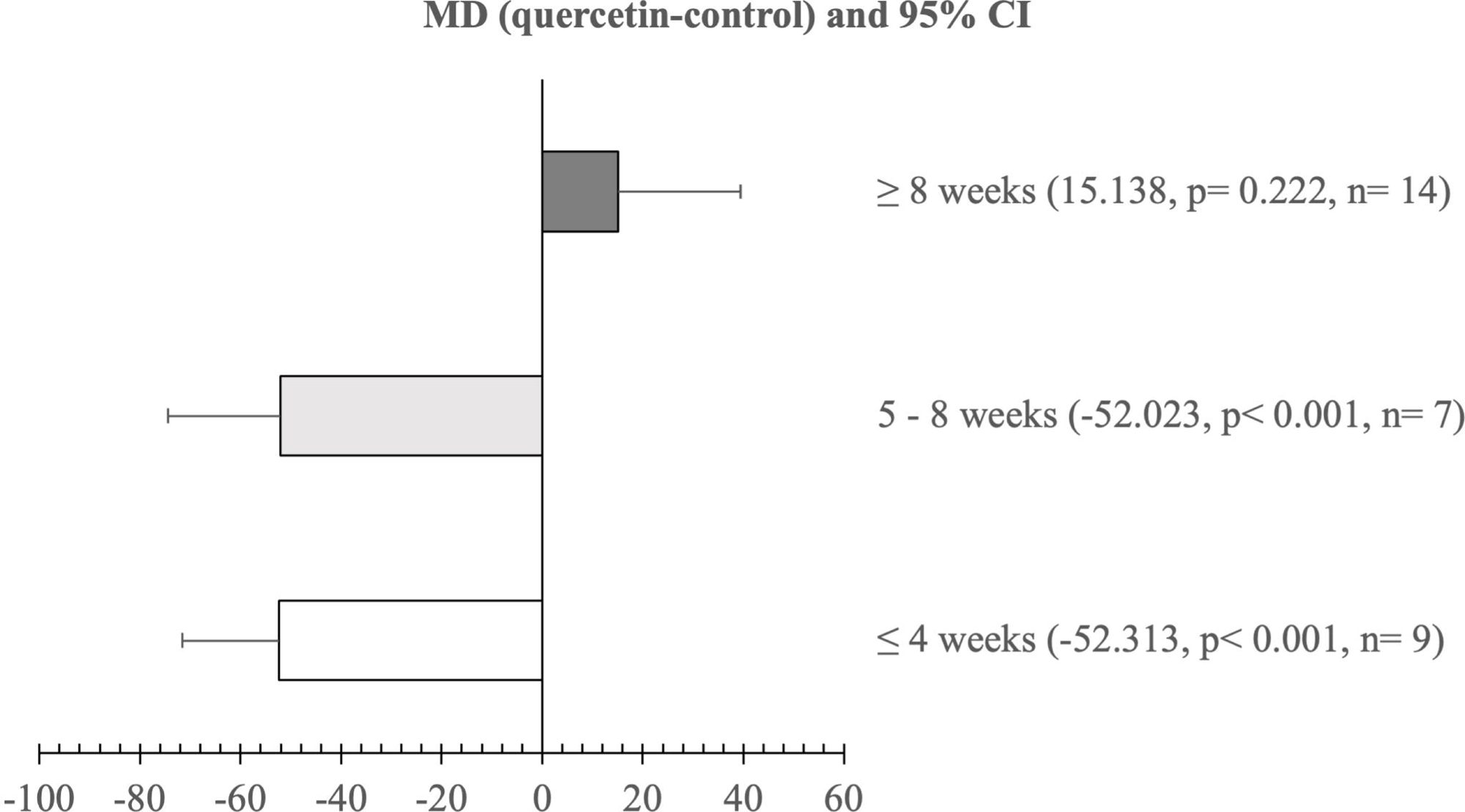
Subgroup analysis for effect of treatment duration on (a) yolk colour, (b) SGPT (U/L), (c) glucose (mg/ dL), and (d) cholesterol (mg/ dL) in laying hens

Figures 5A to C display effect of initial age on LR, ST, and SOD level. Hens aged more than 50 weeks groups had higher (p<0.05) LR and ST than control and less than 50 weeks groups. Moreover, quercetin administration in hens aged less than 50 weeks increased (p<0.05) LR, ST, and SOD level.

**Figure 5.**
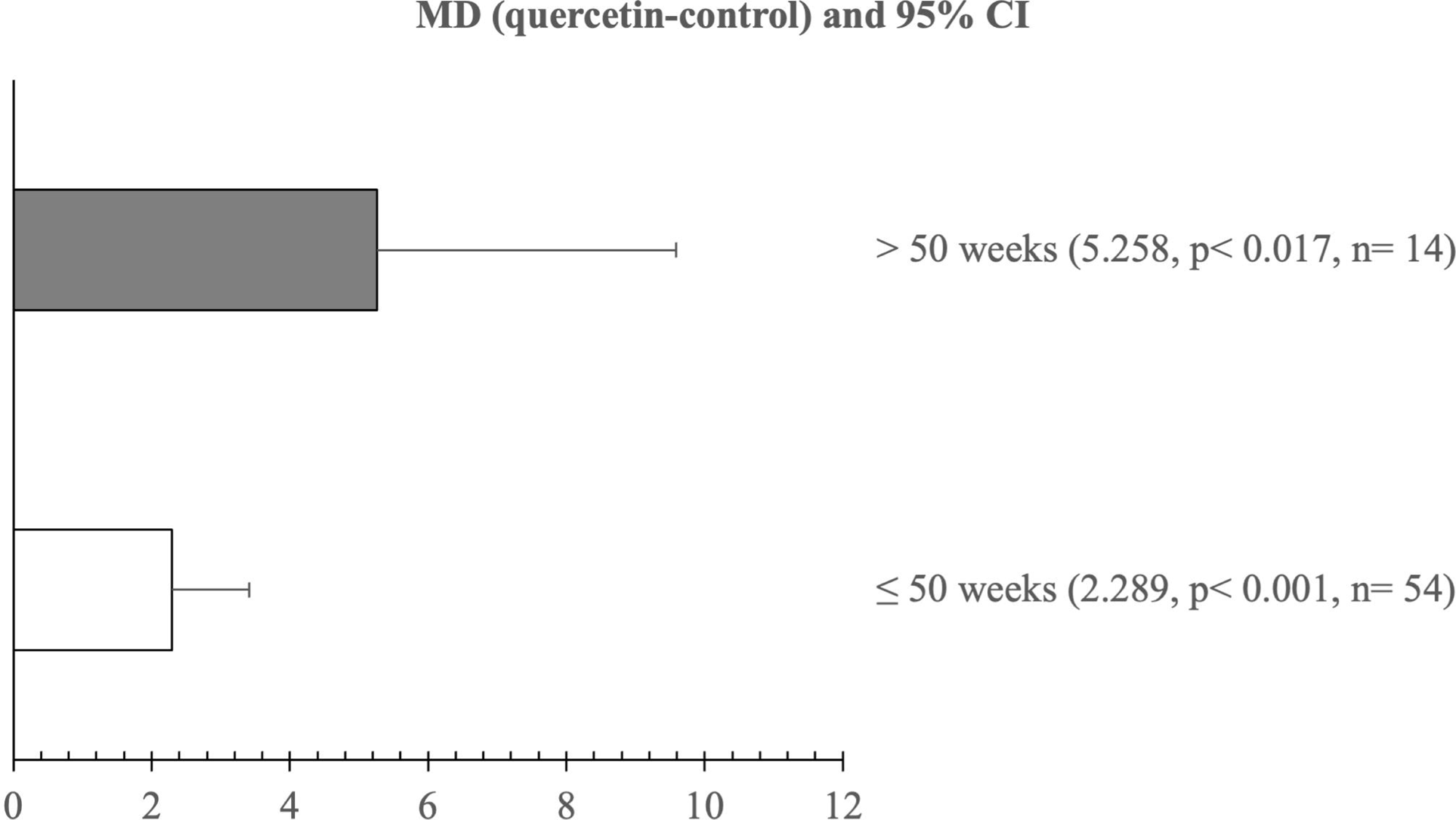
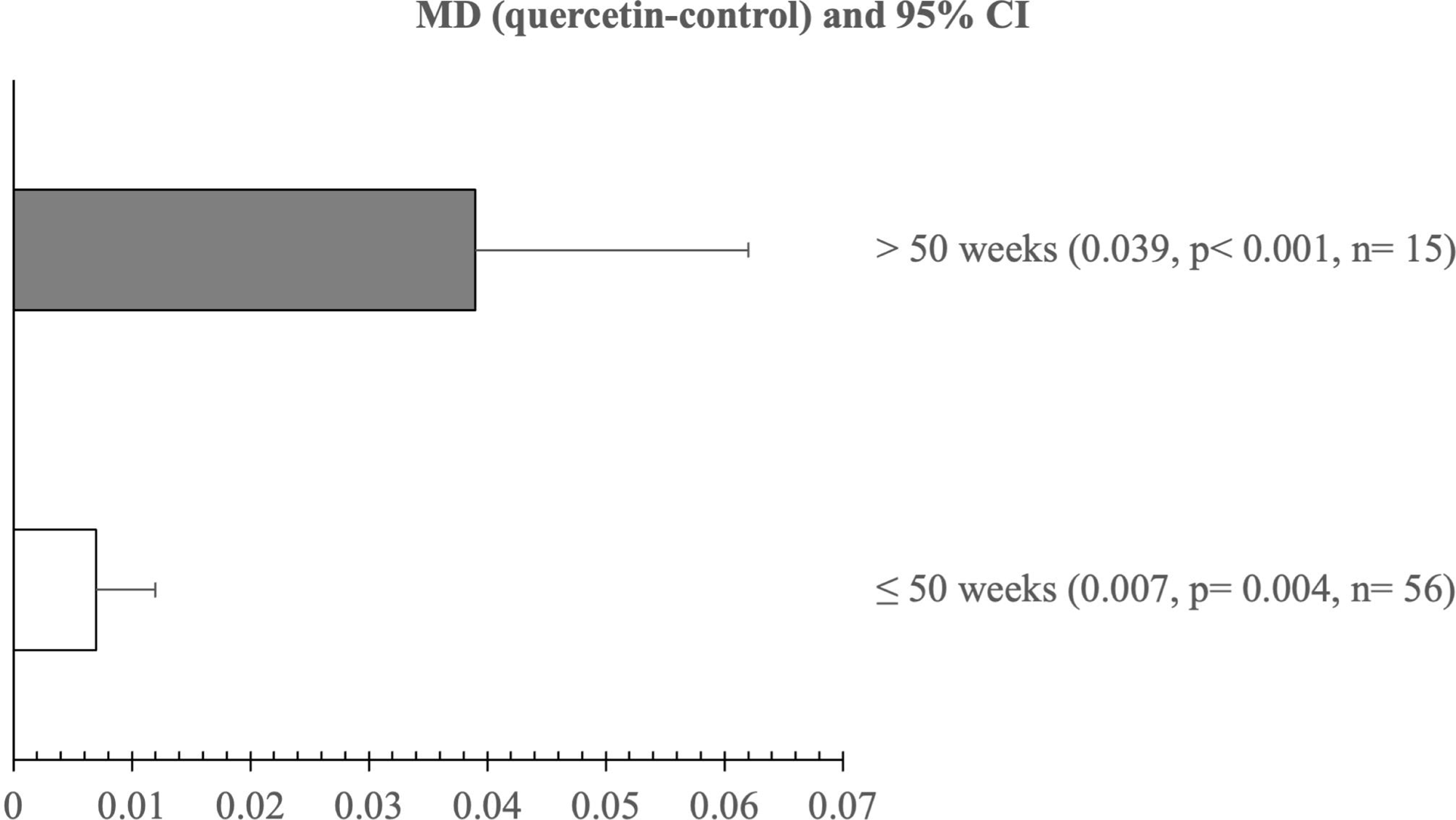
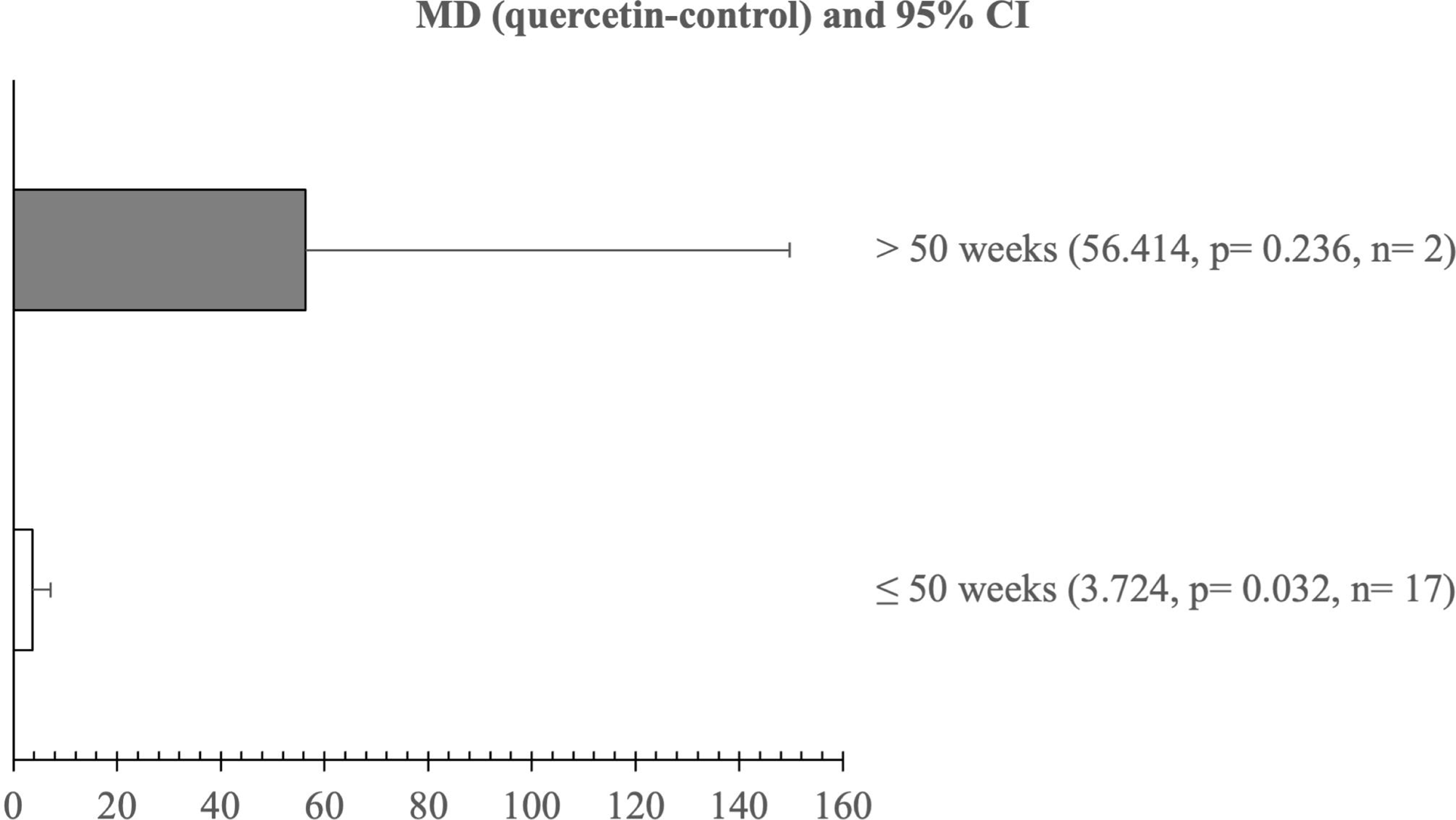
Subgroup analysis of effect of initial age on (a) laying rate (%), (b) shell thickness (mm), and (c) SOD (U/ mL) in laying hens

Form of quercetin influenced LR, FER, YC, cholesterol, and SOD concentration (Figures 6A-E). Extract powder significantly resulted (p<0.05) in highest LR and lowest FER. However, the highest YC and the lowest cholesterol was observed in plant powder. Furthermore, extract powder had no effect on SOD level. In contrast, plant powder improved SOD level.

**Figure 6.**
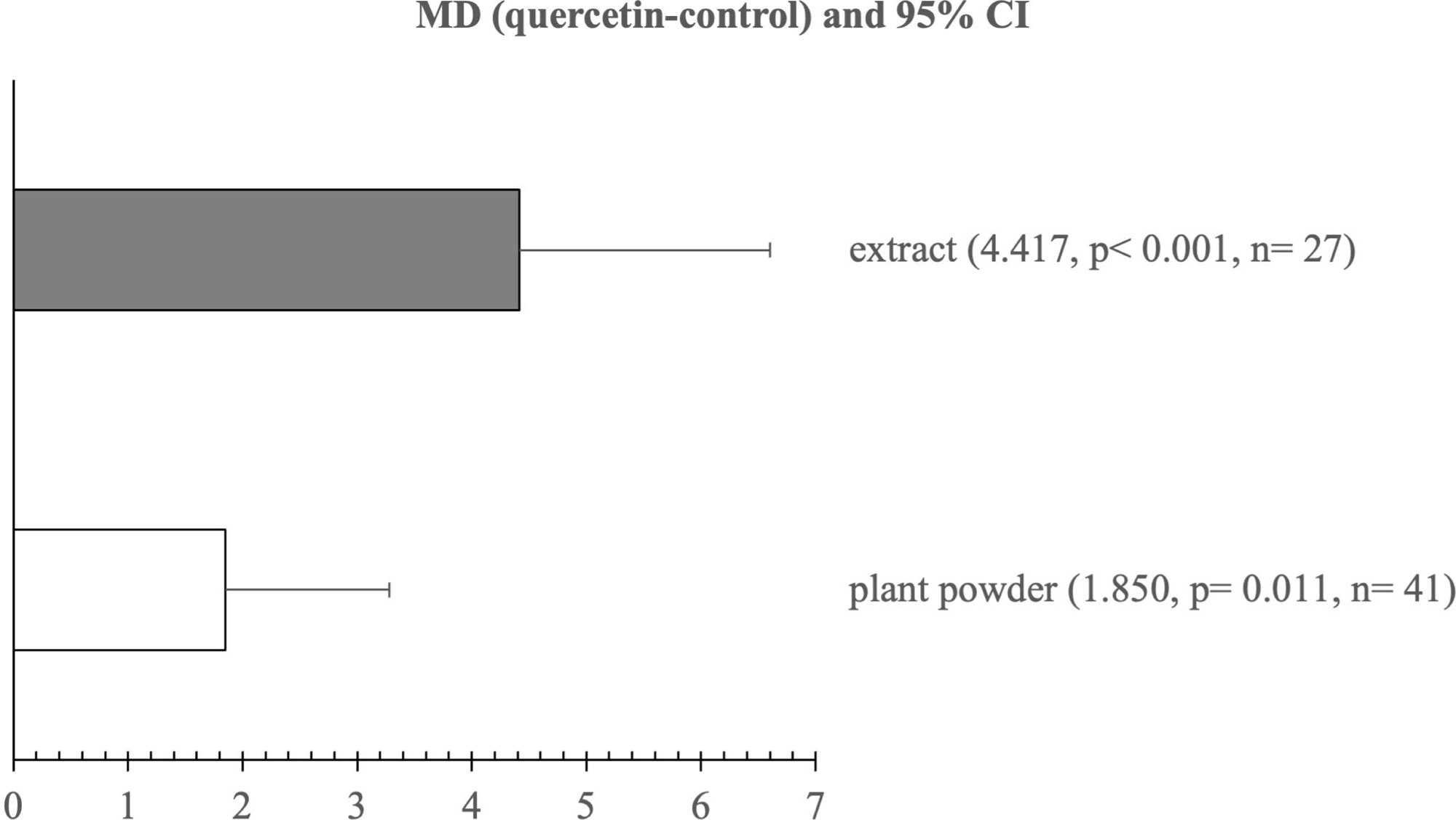
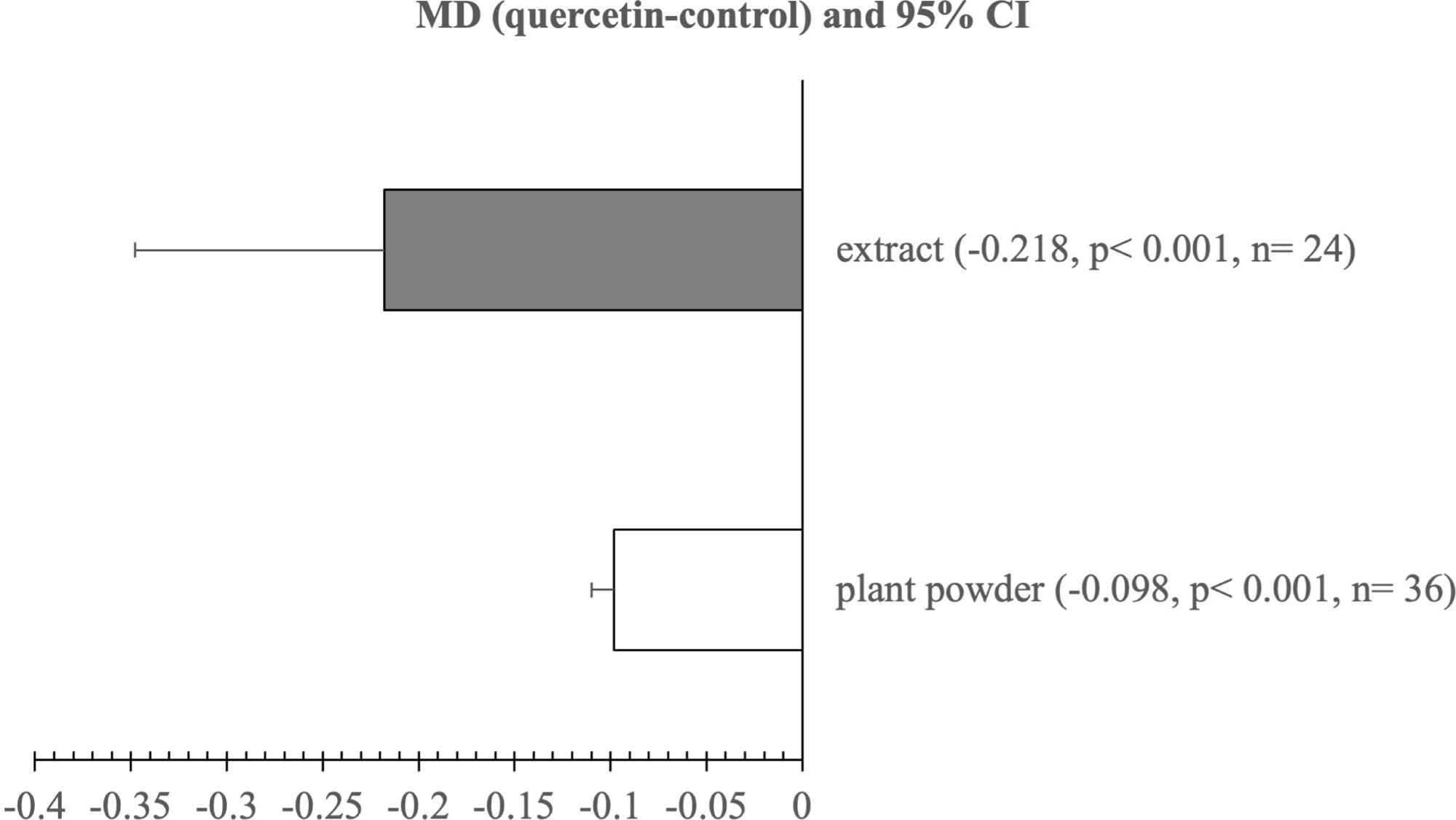
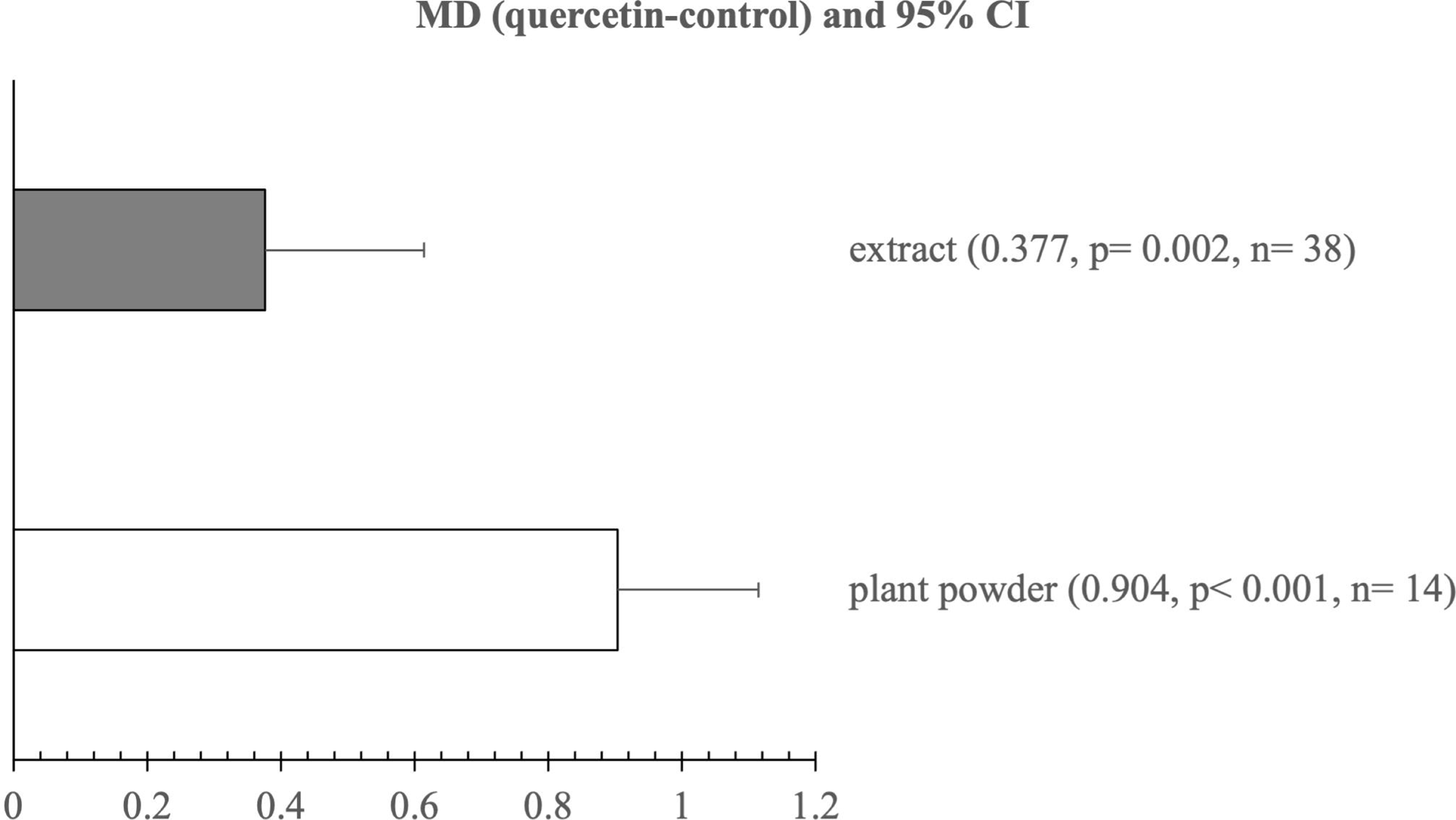
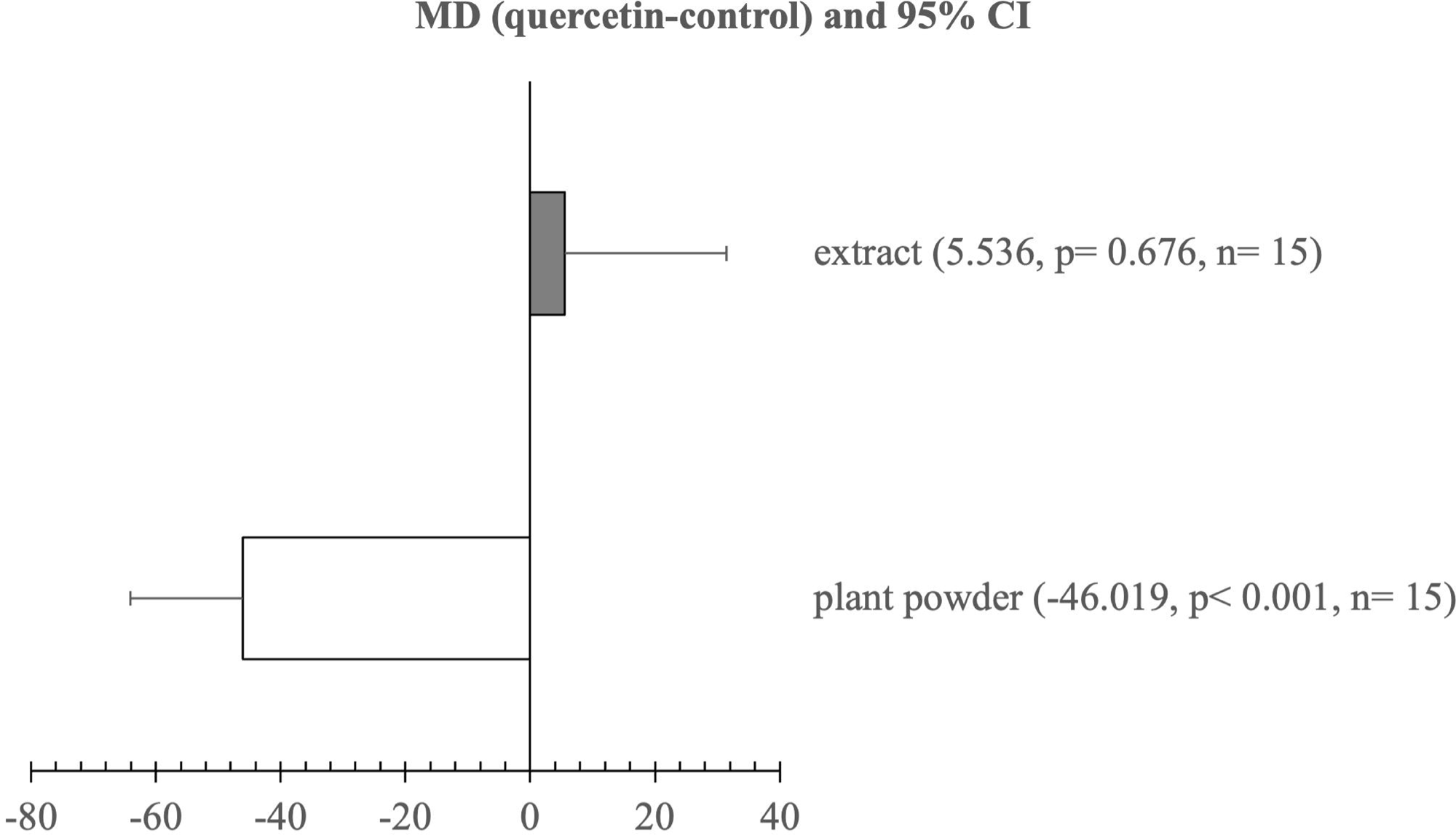
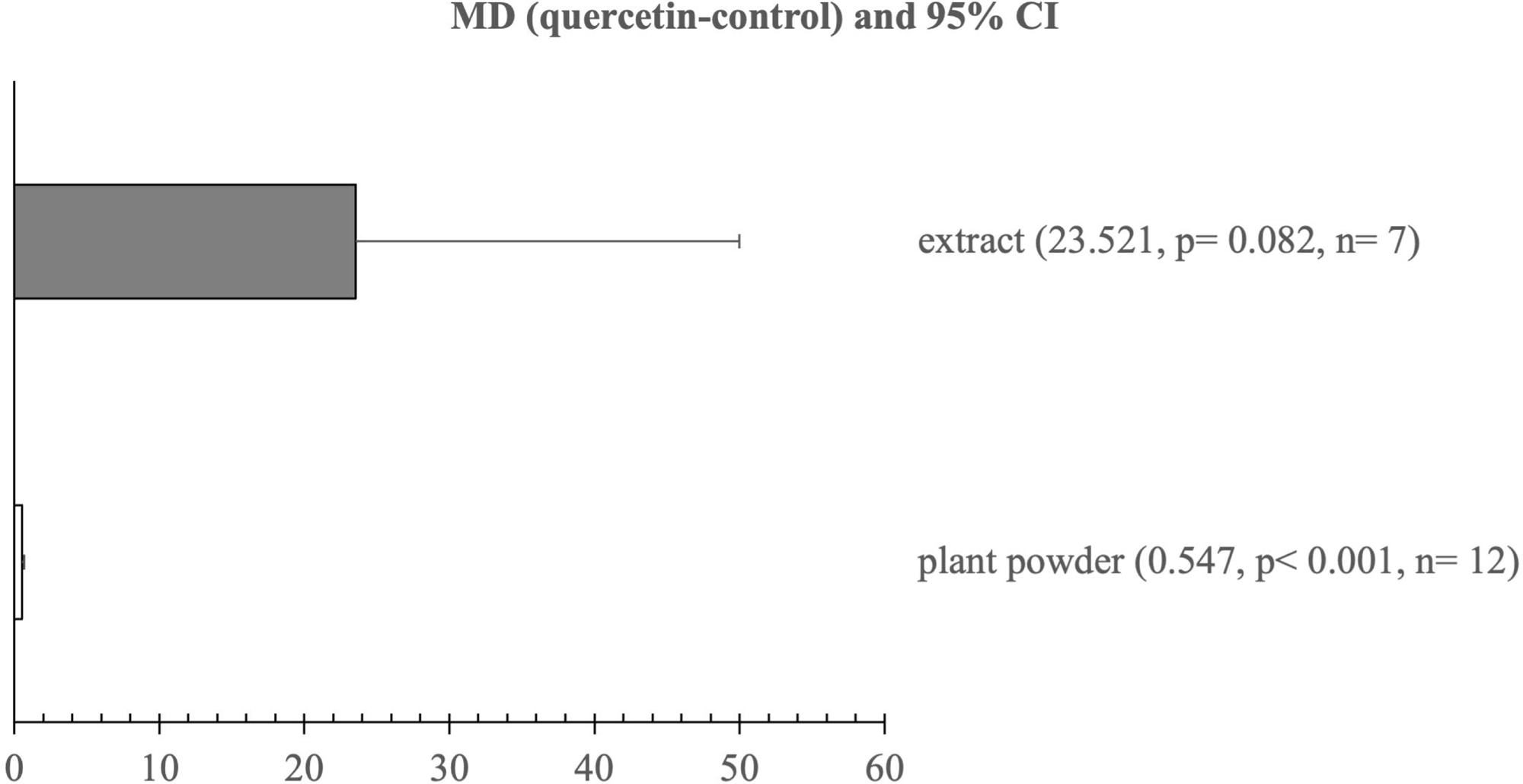
Subgroup analysis for effect of quercetin form on (a) laying rate (%), (b) FER, (c) yolk colour, (d) cholesterol (mg/ dL), and (e) SOD (U/ mL) in laying hens.

### Publication bias

Funnel plots and egger’s test for risk of publication bias are presented in Suppl. Figures 1A to 1P. The LR, FER, EW, HU, ST, YC, SGPT, HDL, LDL, and CAT studies had low (p> 0.05) risk of publication bias. Meanwhile, high risk of publication bias was detected (p<0.05) in glucose, cholesterol, MDA level, and SOD concentration studies.

## DISCUSSION

Quercetin administrations improved performance (FER), production (LR) and egg quality (EW, HU, ST, and YC) indices in the present study. Our findings are similar to Liu et al [7], who reported that quercetin administrations ameliorated LR, EW, FER, HU, ST in laying hens. Moreover, YC score was improved in quercetin-fed hens concomitant with the increased of LR, FER, HU, and ST. Ahmad et al. [10] also demonstrated that quercetin-contained mulberry leaves treatments induced the increase of LR, HU and ST in laying hens. These result might be correlated with the enhanced blood metabolites, antioxidant status, and reproductive hormones levels in quercetin-treated laying hens.

Interestingly, quercetin administration improved blood metabolite in our findings. Total cholesterol and HDL levels in serum are key indicators of lipid metabolism in chickens [47]. Our findings showed that total cholesterol level was decreased and HDL concentration was enhanced in quercetin groups. Our results are in line with study of El-Saadany et al. [12], who found that quercetin treatments decreased cholesterol and increased HDL in serum of laying hens. Similarly, Liu et al. [24] reported that HDL level was elevated and total cholesterol content was reduced in serum of quercetin-fed hens. Lipoprotein produced by liver is crucial for follicle development in hens [48]. The reduced cholesterol and increased HDL contents in serum indicated the improved lipid metabolism and a decreased risk of fatty liver disease in laying hens [49]. Quercetin promotes cholesterol homeostasis by enhancing the selective uptake of HDL-derived lipids [50]. Hepatic function is critically influenced by cholesterol homeostasis condition, whereby imbalance in cholesterol level induce intrahepatic lipid accumulation contributing to liver injury [51]. Moreover, the reduced SGPT level in our study also insisted that quercetin supplementation had hepatoprotective effect in layers. A previous study showed that flavonoid administration lowered SGPT level implying less liver dysfunction [52]. Amevor et al. [22] discovered that quercetin supplementations alleviated hepatic steatosis in aging laying hens. Hence, quercetin administration enhanced hepatic function and modulates lipid metabolism, leading to improved egg production and quality in laying hens.

Our study showed that the anti-oxidant activity was increased in quercetin-supplemented hens as indicated by the enhance of SOD concentration and the reduce of MDA level. The result is similar to study of Liu et al. [14], whereby MDA level was declined and SOD concentration was elevated in serum of the quercetin-supplemented hens. Interestingly, similar results are also found in aging [22] and heat-stressed laying hens [21]. A study of Lin et al. [17] also demonstrated that quercetin-contained mulberry leaves treatment had increased SOD level in line with the reduced MDA in serum of laying hens. The SOD, an antioxidant enzyme, is a vital first line defence system against oxidative stress agent [53]. It has also significant role for ovarian development [54]. On the other hand, MDA level is significantly indicator for oxidative stress in serum and tissue [55]. Our results implied that quercetin dietary alleviated oxidative stress by enhancing anti-oxidant defence. The reduced oxidative stress contributed to enhancement in the performance, production, and egg quality in laying hens.

Moreover, previous studies reported that quercetin treatments affected reproductive hormone levels, including oestradiol (E2) and follicle-stimulating hormone (FSH). Quercetin-treated hens had higher E2 level, a key oestrogen produced by ovarian follicles [22]. Yang et al. [11] demonstrated that quercetin dietary increased significantly E2 level concomitant with enhanced ovarian follicle numbers, which is associated with improved laying performance. E2 content also affects egg quality, especially shell thickness in laying hens. Wistedt et al. [56] reported that E2 treatment improved shell thickness in laying hens. Moreover, high FSH level was found in quercetin-administrated hens [57]. The study also found that FSH treatment promoted follicle development with increased yolk deposition, angiogenesis, and cell proliferation. High FSH content was in line with the increase of laying rate in hens [58]. Thus, quercetin treatment effectively optimized reproductive hormone contents promoting the increase of egg production and egg quality.

### Limitations

The present study solely synthesized effect of a single quercetin treatment. Combinations of quercetin with other antioxidants, enzymes, vitamins, and minerals were excluded. Breed-based subgroup analysis was no performed because of insufficient studies for each breed.

## CONCLUSION

Quercetin treatment effectively improved performance, egg production, egg quality, blood metabolites, and anti-oxidant defence system. Quercetin dose linearly impacted the SGPT, glucose, and cholesterol contents. However, quercetin dose had quadratic effect on LR, FER, and SOD content. The effective quercetin dose for optimizing LER, FER, and SOD content was 400 to 600 mg/kg. Supplementation of quercetin for more than 5 and 8 weeks improved YC. Moreover, quercetin administration for less than 8 enhanced blood metabolites. Quercetin effectively elevated LR and ST in all aged hens. Quercetin dietary in extract powder form enhanced egg production, whereas quercetin supplementation in plant powder increased egg quality.

## Supporting information

https://drive.google.com/file/d/11GqfKO4qGirbqfAuaZSMFEChveiSJ2HZ/view?usp=sharing

## ACKNOWLEDGMENTS

Authors would like to thank National Research and Innovation Agency for providing comprehensive access to literature databases (Scopus and Web of Science). This research has no grant from any funding agency.

## REFERENCES

1. Mottet A, Tempio G. Global poultry production: current state and future outlook and challenges. J World’s Poult Sci. 2017;73(2):245–56. 10.1017/S0043933917000071

2. Wasti S, Sah N, Mishra B. Impact of heat stress on poultry health and performances, and potential mitigation strategies. Animals. 2020;10(8):1266. 10.3390/ani10081266

3. Wang J, Yue H, Wu S, Zhang H, Qi G. Nutritional modulation of health, egg quality and environmental pollution of the layers. Anim Nutr. 2017;3(2):91–6. 10.1016/j.aninu.2017.03.001

4. Wallace RJ, Oleszek W, Franz C, Hahn I, Baser KHC, Mathe A, et al. Dietary plant bioactives for poultry health and productivity. Br Poult Sci. 2010;51(4):461–87. 10.1080/00071668.2010.506908

5. Aghababaei F, Hadidi M. Recent advances in potential health benefits of quercetin. Pharmaceuticals. 2023;16(7):1020. 10.1080/10.3390/ph16071020

6. Hu T, Yue J, Tang Q, Cheng KW, Chen F, Peng M, et al. The effect of quercetin on diabetic nephropathy (DN): a systematic review and meta-analysis of animal studies. Food Funct. 2022;13(9):4789–803. 10.1080/10.1039/D1FO03958J

7. Liu Y, Li Y, Liu HN, Suo YL, Hu LL, Feng XA, et al. Effect of quercetin on performance and egg quality during the late laying period of hens. Brit Poult Sci. 2013;54(4):510–4. 10.1080/00071668.2013.799758

8. Simitzis P, Spanou D, Glastra N, Goliomytis M. Impact of dietary quercetin on laying hen performance, egg quality and yolk oxidative stability. Anim Feed Sci Technol. 2018;239:27–32. 10.1016/j.anifeedsci.2018.03.004

9. Amevor FK, Cui Z, Ning Z, Du X, Jin N, Shu G, et al. Synergistic effects of quercetin and vitamin E on egg production, egg quality, and immunity in aging breeder hens. Poult Sci. 2021;100(12):101481. 10.1016/j.psj.2021.101481

10. Ahmad S, Khalique A, Pasha T, Mehmood S, Ahmad SS, Khan A, et al. Influence of moringa oleifera leaf meal used as phytogenic feed additive on the serum metabolites and egg bioactive compounds in commercial layers. Braz J Poult Sci. 2018;20(2):325–32. 10.1590/1806-9061-2017-0606

11. Yang JX, Chaudhry MT, Yao JY, Wang SN, Zhou B, Wang M, et al. Effects of phyto_oestrogen quercetin on productive performance, hormones, reproductive organs and apoptotic genes in laying hens. J Anim Physiol Anim Nutr. 2018;102(2):505–13. 10.1111/jpn.12778

12. El-Saadany A, El-Barbary AM, El-Salam AA, Ahmed MM, Shreif EY. Nutritional and physiological evaluation of quercetinas a phytogenic feed additive in laying hens. J Anim Feed Sci. 2022;31(3):249–57. 10.22358/jafs/150080/2022

13. Shen M, Li T, Lu J, Qu L, Wang K, Hou Q, et al. Effects of supplementation of Moringa Oleifera leaf powder on some reproductive performance in laying hens. Braz J Poult Sci. 2022;24(2):eRBCA-2021-1537. 10.1590/1806-9061-2021-1537

14. Liu J, Fu Y, Zhou S, Zhao P, Zhao J, Yang Q, et al. Comparison of the effect of quercetin and daidzein on production performance, anti-oxidation, hormones, and cecal microflora in laying hens during the late laying period. Poult Sci. 2023;102(6):102674. 10.1016/j.psj.2023.102674

15. Fu Y, Zhou J, Schroyen M, Lin J, Zhang H, Wu S, et al. Dietary supplementation with calcitriol or quercetin improved eggshell and bone quality by modulating calcium metabolism. Anim Nutr. 2024;18:340–55. 10.1016/j.aninu.2024.04.007

16. Ahmad S, Khalique A, Pasha TN, Mehmood S, Hussain K, Ahmad S, et al. Effect of *Moringa oleifera* (Lam.) pods as feed additive on egg antioxidants, chemical composition and performance of commercial layers. SA J An Sci. 2017;47(6):864. 10.4314/sajas.v47i6.14

17. Lin WC, Lee MT, Chang SC, Chang YL, Shih CH, Yu B, et al. Effects of mulberry leaves on production performance and the potential modulation of antioxidative status in laying hens. Poult Sci. 2017;96(5):1191–203. 10.3382/ps/pew350

18. Su BW, Lin WC, Lin LJ, Huang CM, Chuang WY, Wu DJ, et al. Laying diet supplementation with *Ricinus communis* L. leaves and evaluation of productive performance and potential modulation of antioxidative status. J Poult Sci. 2020;57(4):259–69. 10.2141/jpsa.0190077

19. Wei Y, Liu Y, Li G, Guo Y, Zhang B. Effects of quercetin and genistein on egg quality, lipid profiles, and immunity in laying hens. J Sci Food Agric. 2024;104(1):207–14. 10.1002/jsfa.12910

20. Abid AR, Ahmed SK. Egg quality of hen affected by different levels of quercetin. Biochem Cell Arch. 2019;19(2):2823–30. 10.35124/bca.2019.19.2.2823

21. Cao X, Amevor FK, Du X, Wu Y, Xu D, Wei S, et al. Supplementation of the combination of quercetin and vitamin E alleviates the effects of heat stress on the uterine function and hormone synthesis in laying hens. Animals. 2024;14(11):1554. 10.3390/ani14111554

22. Amevor FK, Cui Z, Du X, Ning Z, Shu G, Jin N, et al. Combination of quercetin and vitamin E supplementation promotes yolk precursor synthesis and follicle development in aging breeder hens via liver–blood–ovary signal axis. Animals. 2021;11(7):1915. 10.3390/ani11071915

23. Iskender H, Yenice G, Dokumacioglu E, Kaynar O, Hayirli A, Kaya A. The effects of dietary flavonoid supplementation on the antioxidant status of laying hens. Rev Bras Cienc Avic. 2016;18(4):663–8. 10.1590/1806-9061-2016-0356

24. Liu J, Liu J, Zhou S, Fu Y, Yang Q, Li Y. Effects of quercetin and daidzein on egg quality, lipid metabolism, and cecal short-chain fatty acids in layers. Front Vet Sci. 2023;10:1301542. 10.3389/fvets.2023.1301542

25. Damaziak K, Riedel J, Gozdowski D, Niemiec J, Siennicka A, Róg D. Productive performance and egg quality of laying hens fed diets supplemented with garlic and onion extracts. J Appl Poult Res. 2017;26(3):337–49. 10.3382/japr/pfx001

26. Huang Z, Dai H, Jiang J, Ye N, Zhu S, Wei Q, et al. Dietary mulberry-leaf flavonoids improve the eggshell quality of aged breeder hens. Theriogenology. 2022;179:177–86. 10.1016/j.theriogenology.2021.11.019

27. Abid AR, Ahmed SK. Influence of quercetin on some physiological measurements of layer hens. Plant Arch. 2019;19(2):3575–82.

28. Whiting IM, Pirgozliev V, Kljak K, Orczewska-Dudek S, Mansbridge SC, Rose SP, et al. Feeding dihydroquercetin in wheat-based diets to laying hens: impact on egg production and quality of fresh and stored eggs. Brit Poult Sci. 2022;63(6):735–41. 10.1080/00071668.2022.2090229

29. Amevor FK, Cui Z, Du X, Ning Z, Deng X, Xu D, et al. Synergy between dietary quercetin and vitamin E supplementation in aged hen’s diet improves hatching traits, embryo quality, and antioxidant capacity of chicks hatched from eggs subjected to prolonged storage. Front Physiol. 2022;13:873551. 10.3389/fphys.2022.873551

30. Amevor FK, Cui Z, Du X, Feng J, Shu G, Ning Z, et al. Synergy of dietary quercetin and vitamin E improves cecal microbiota and its metabolite profile in aged breeder hens. Front Microbiol. 2022;13:851459. 10.3389/fmicb.2022.851459

31. Liu HN, Liu Y, Hu LL, Suo YL, Zhang L, Jin F, et al. Effects of dietary supplementation of quercetin on performance, egg quality, cecal microflora populations, and antioxidant status in laying hens. Poult Sci. 2014;93(2):347–53. 10.3382/ps.2013-03225

32. Iskender H, Yenice G, Dokumacioglu E, Kaynar O, Hayirli A, Kaya A. Comparison of the effects of dietary supplementation of flavonoids on laying hen performance, egg quality and egg nutrient profile. Brit Poult Sci. 2017;58(5):550–6. 10.1080/00071668.2017.1349297

33. Ying Y, Chun-yan H, Tabassum CM, Ling L, Jia-ying Y, Sheng-nan W, et al. Effect of quercetin on egg quality and components in laying hens of different weeks. J Northeast Agric Univ. 2015;22(4):23–32. 10.1016/S1006-8104(16)30015-0

34. Wang R. Combining Data from Multiple Studies. NEJM Evid. 2023;2(5):1–2. 10.1056/EVIDe2300066

35. Jain S. Meta-Analysis: A higher quality of evidence in clinical research pyramid. Int J Sci Res. 2018;9(3):340–9.

36. Higgins JPT, Thomas J, Chandler J, Chumpston M, Li T, Page MJ, et al. Cochrane Handbook for Systematic Reviews of Interventions Version 6.4. In: Cochrane Handbook for Systematic Reviews of Interventions Version 64 [Internet]. London, United Kingdom: Cochrane; 2023 [cited 2024 Oct 24]. (Version 6.4.). Available from:https://www.training.cochrane.org/handbook

37. Rohatgi A. WebPlotDigitizer [Internet]. California, USA; 2024 [cited 2024 Oct 24]. Available from: https://automeris.io

38. Ranga Niroshan Appuhamy JAD, Strathe AB, Jayasundara S, Wagner-Riddle C, Dijkstra J, France J, et al. Anti-methanogenic effects of monensin in dairy and beef cattle: A meta-analysis. J Dairy Sci. 2013;96(8):5161–73. 10.3168/jds.2012-5923

39. Higgins JPT. Measuring inconsistency in meta-analyses. BMJ. 2003;327(7414):557–60. 10.1136/bmj.327.7414.557

40. Egger M, Smith GD, Schneider M, Minder C. Bias in meta-analysis detected by a simple, graphical test. BMJ. 1997;315(7109):629–34. 10.1136/bmj.315.7109.629

41. Budiyanto A, Hartanto S, Widayanti R, Kurnianto H, Wardi W, Haryanto B, et al. Impact of melatonin administration on sperm quality, steroid hormone levels, and testicular blood flow parameters in small ruminants: A meta-analysis. Vet World. 2024;911–21. 10.14202/vetworld.2024.911-921

42. St-Pierre NR. Invited Review: Integrating Quantitative Findings from Multiple Studies Using Mixed Model Methodology. J Dairy Sci. 2001;84(4):741–55. 10.3168/jds.S0022-0302(01)74530-4

43. R Core Team. R: A Language and Environment for Statistical Computing. Vienna, Austria: R Foundation for Statistical Computing; 2023.

44. Viechtbauer W. Conducting Meta-Analyses in *R* with the **metafor** Package. J Stat Soft [Internet]. 2010 [cited 2025 Feb 23];36(3). Available from: http://www.jstatsoft.org/v36/i03/

45. Wickham H. ggplot2: elegant graphics for data analysis. Second edition. Switzerland: Springer; 2016. 260 p. (Use R!).

46. Microsoft Corporation. Microsoft Excel [Internet]. 2019. Available from: https://office.microsoft.com/excel

47. Zaefarian F, Abdollahi MR, Cowieson A, Ravindran V. Avian Liver: The Forgotten Organ. Animals. 2019;9(2):63. 10.3390/ani9020063

48. Van Eck LM, Enting H, Carvalhido IJ, Chen H, Kwakkel RP. Lipid metabolism and body composition in long-term producing hens. Worlds Poult Sci J. 2023;79(2):243–64. 10.1080/00439339.2023.2189206

49. Attia YA, Al Sagan AA, Hussein E sayed OS, Olal MJ, Ebeid TA, Alhotan RA, et al. Antioxidant status, lipid metabolism, egg fatty acids, and nutritional index of white-egg laying hens fed flaxseed cake. J Poult Sci. 2024;61(0):1–13. 10.2141/jpsa.2024010

50. Ren K, Jiang T, Zhao GJ. Quercetin induces the selective uptake of HDL-cholesterol via promoting SR-BI expression and the activation of the PPARγ/LXRα pathway - Food & Function (RSC Publishing). Food Funct. 2018;9:624–35. 10.1039/C7FO01107E

51. Duan Y, Gong K, Xu S, Zhang F, Meng X, Han J. Regulation of cholesterol homeostasis in health and diseases: from mechanisms to targeted therapeutics. Sig Transduct Target Ther. 2022;7(1):265. 10.1038/s41392-022-01125-5

52. Kuttiappan A, Chenchula S, Vanangamudi M, Bhatt S, Chikatipalli R, Shaila Bhanu P, et al. Hepatoprotective effect of flavonoid rich fraction of Sesbania grandiflora: Results of In vivo, in vitro, and molecular docking studies. J Ayurveda Integr Med. 2024;15(5):101036. 10.1016/j.jaim.2024.101036

53. Ighodaro OM, Akinloye OA. First line defence antioxidants-superoxide dismutase (SOD), catalase (CAT) and glutathione peroxidase (GPX): Their fundamental role in the entire antioxidant defence grid. Alex J Med. 2018;54(4):287–93. 10.1016/j.ajme.2017.09.001

54. Zheng M, Liu Y, Zhang G, Yang Z, Xu W, Chen Q. The applications and mechanisms of superoxide dismutase in medicine, food, and cosmetics. Antioxidants. 2023;12(9):1675. 10.3390/antiox12091675

55. Cordiano R, Di Gioacchino M, Mangifesta R, Panzera C, Gangemi S, Minciullo PL. Malondialdehyde as a potential oxidative stress marker for allergy-oriented diseases: An update. Molecules. 2023;28(16):5979. 10.3390/molecules28165979

56. Wistedt A, Ridderstråle Y, Wall H, Holm L. Exogenous estradiol improves shell strength in laying hens at the end of the laying period. Acta Vet Scand. 2014;56(1):34. 10.1186/1751-0147-56-34

57. Ma Y, Yao J, Zhou S, Mi Y, Tan X, Zhang C. Enhancing effect of FSH on follicular development through yolk formation and deposition in the low-yield laying chickens. Theriogenology. 2020;157:418–30. 10.1016/j.theriogenology.2020.07.012

58. Prastiya RA, Madyawati SP, Sari SY, Nugroho AP. Effect of follicle-stimulating hormone and luteinizing hormone levels on egg-laying frequency in hens. Vet World. 2022;15:2890–5. 10.14202/vetworld.2022.2890-2895

