## Supplementary figures and images for "Quercetin treatments alleviate production, egg quality, blood metabolites, and anti-oxidant defence in laying hens : A meta-analysis study"

### https://drive.google.com/file/d/11GqfKO4qGirbqfAuaZSMFEChveiSJ2HZ/view?usp=sharing

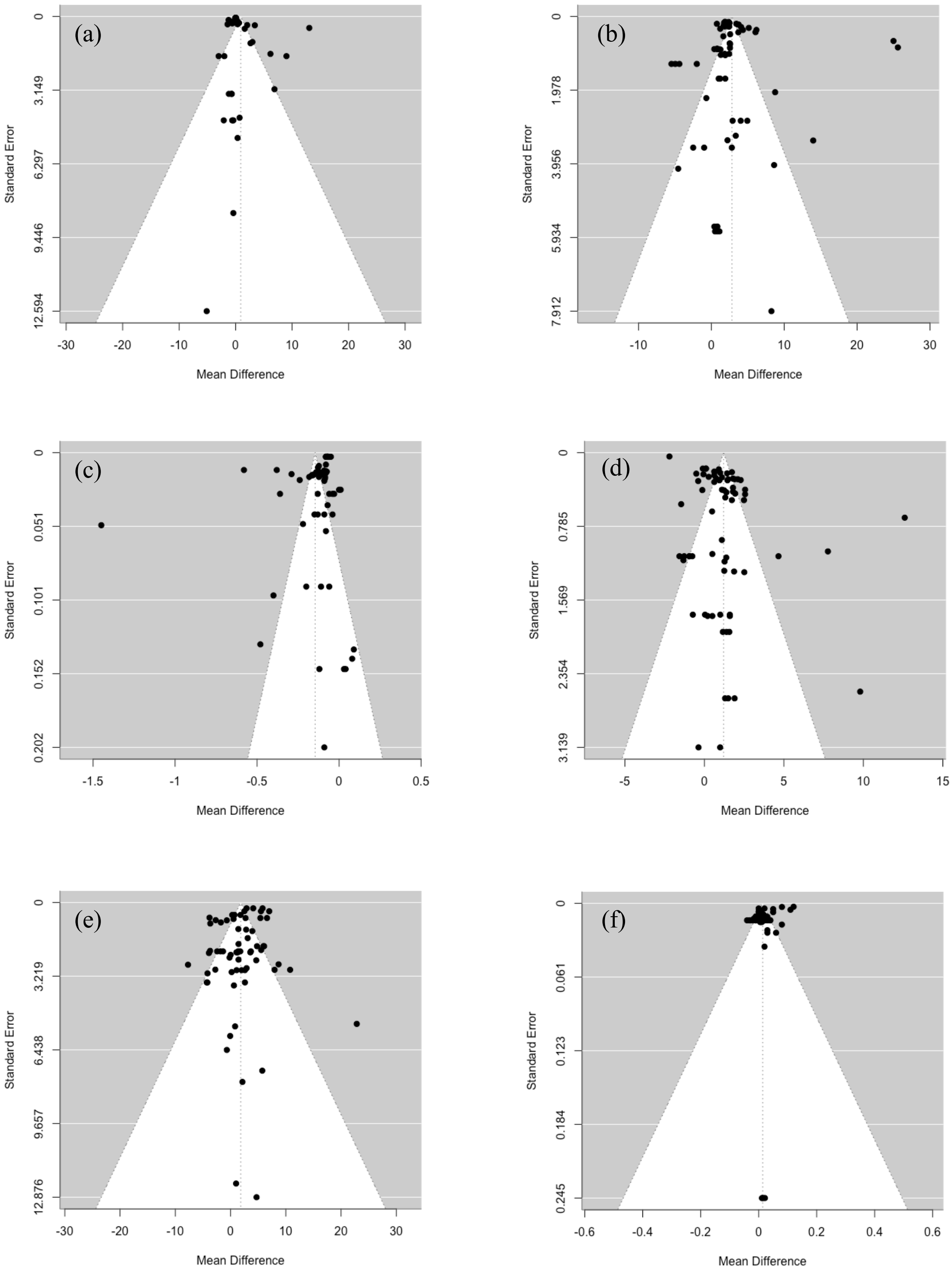

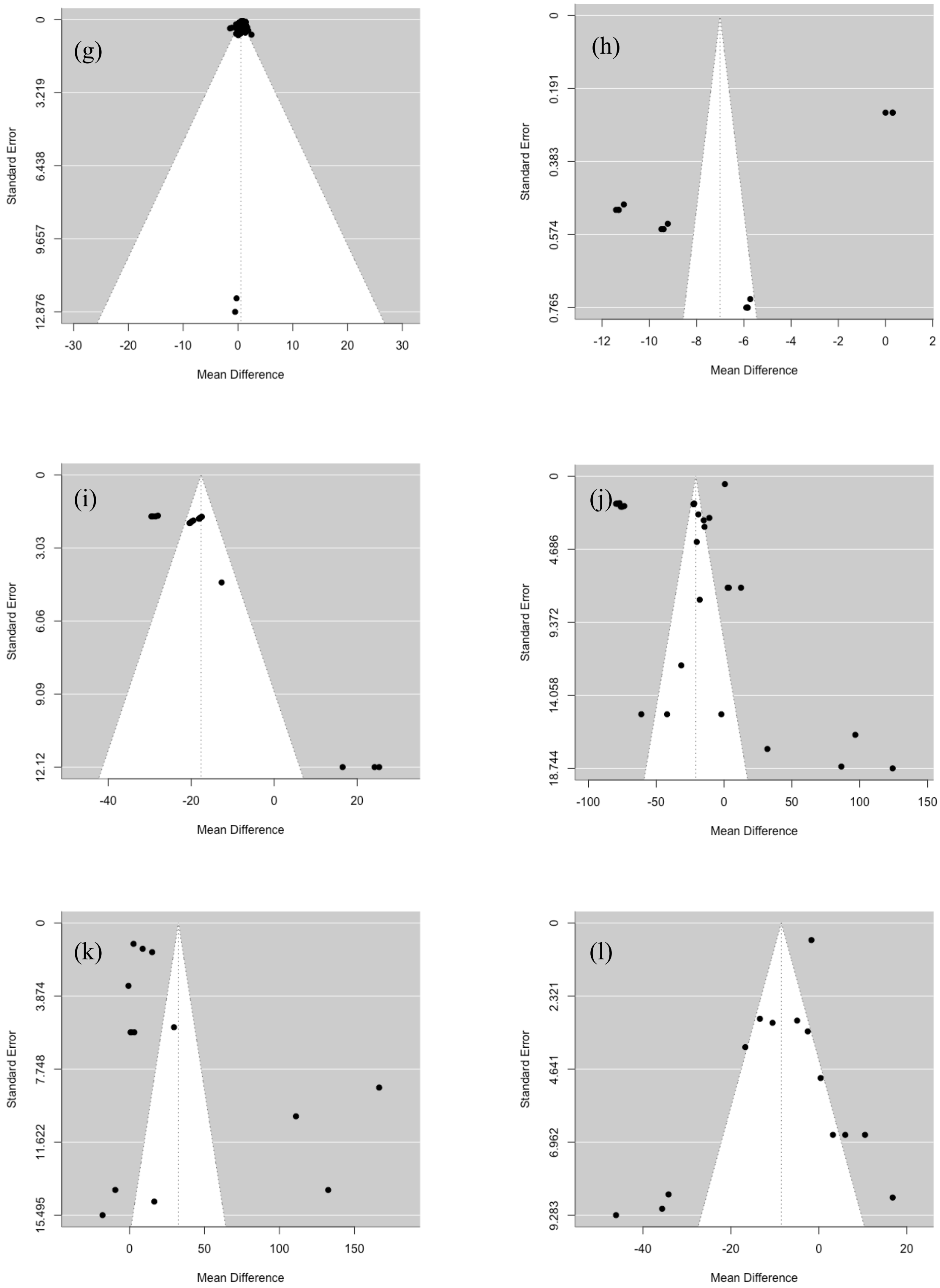

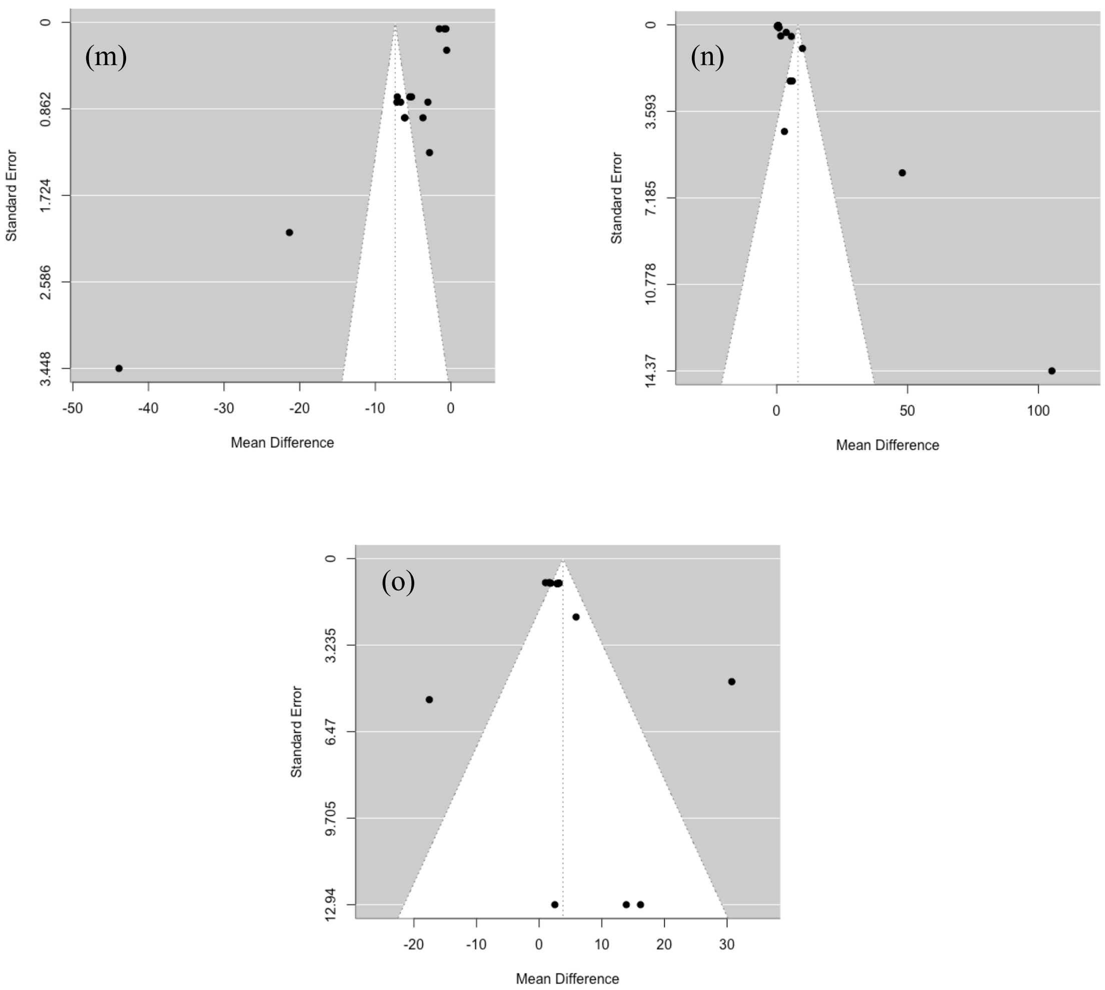
